# Pre-plasma cells proliferate and differentiate at the germinal center-medulla interface of antigen-draining lymph nodes

**DOI:** 10.1101/2024.07.26.605240

**Authors:** Laurine Binet, Lucas Bremond, Marius Duparc, Chuang Dong, Léa David, Noudjoud Attaf, Laurine Gil, Matthieu Fallet, Thomas Boudier, Bertrand Escalière, Lionel Chasson, Myriam Moussa, Lucas Bertoia, Samuel Origlio, Sylvain Bigot, Carole Siret, Serge Van de Pavert, Jean-Marc Navarro, Mauro Gaya, Pierre Milpied

## Abstract

High affinity antibody-producing plasma cells (PC) generated in germinal centers (GC) are crucial for durable immunity after vaccination or infection. The selection of high affinity B cells in the GC light zone instructs PC differentiation in a subset of cells, but the cellular transitions and spatial organization of GC to PC differentiation remain poorly understood. Here, we have used a mouse model to track GC-derived B cells with integrative single-cell and spatial analyses in draining lymph node after immunization or infection. We first identified putative PrePC cells in scRNA-seq datasets, then enriched those cells through their specific surface phenotype for further analysis of their gene expression trajectories and BCR repertoire. We found a continuum of actively proliferating transitional states bridging selected LZ GC B cells and recently exported PCs, with gradually increasing levels of endoplasmic reticulum stress-associated genes and Ig transcripts. Spatial analyses revealed that recently differentiated PC continued their maturation and affinity-restricted proliferation at a previously uncharacterized interface between the DZ and extensions of the lymph node medulla. Our findings provide insights into the intermediate stages and microenvironmental factors involved in the differentiation of GC B cells into PC, with implications for vaccine development and understanding antibody responses.

## INTRODUCTION

B cell immune responses are important for long term protection against all kinds of pathogens, after natural infection or vaccination. Germinal Centers (GCs), micro-anatomical structures that form within B cell follicles in secondary lymphoid organs after T cell-dependent B cell activation, play a crucial part for long term antibody-based immunity. In GCs, the affinity maturation cyclic process enables the diversification and overall gain in affinity of antigen-specific antibodies expressed as surface B cell receptors (BCR) by GC B cells. In a maturation cycle, B cells first undergo cell division and somatic hypermutation (SHM) in the dark zone (DZ) in order to diversify their BCR before migrating to the light zone (LZ) and being selected based on the affinity of their BCR. Most studies have shown that selection broadly follows the affinity-dependent selection model^1^. In this model, the really low affinity B cells die by apoptosis, the low affinity ones differentiate in memory B cells (MBC), the intermediate ones recycle in the DZ and the high affinity ones differentiate in PC^1–4^. However, that model, which has been established in clonally-restricted systems using simple model antigens, has been challenged by observations where a broad range of affinities can be detected among GC-derived PC in response to infection or immunization with complex antigens in polyclonal systems^5,6^.

The differentiation of GC-derived MBC and plasma cells (PC) results in diversified and affinity-enhanced antibody specificities being expressed in long-lived cell types for long-term immune protection. In that regard, GC-derived PC have been shown to be the primary source of long-lived high-affinity antibody producing cells that home to the bone marrow^7^. The differentiation of GC B cells into long-lived PC results from the induction of gene expression modifications through different signals. First, GC B cells have to internalize the antigen contained in immune complexes at the FDC membrane before they present it to T follicular helper cells (T_FH_) on MHC class II molecules and receive co-stimulatory signals. The BCR signal is also important in itself thanks to the repression of *Bcl6* transcription^8,9^, responsible for the GC profile, and the induction of *Irf4* expression^10,11^. The BCR affinity allows to be more competitive and to extract more antigen at the FDC surface, which has been correlated with a better help from T_FH_ cells ^12–15^. These cells allow the selection of GC B cells through a competitive access to co-stimulatory signals such as CD40L and IL21. CD40 signaling reinforces BCR signaling through the blockade of *Bcl6* expression and the induction of *Irf4* in a dose-dependent manner^16–18^. Finally, IL21 seems to have a dual role in the GC. It induces the expression of Bcl6, but when it synergizes with the BCR signal and the CD40 signal, it allows the expression of Irf4 and Blimp1 (encoded by *Prdm1*), two markers important for terminal PC differentiation^19–22^.

The precise mapping of stepwise processes for PC differentiation has been mostly studied in *ex vivo* differentiation assays starting from naïve B cells or MBC. Although the situation may differ for the GC to PC differentiation *in vivo*, those assays have generated important findings at the genetic level. Notably, because some GC markers such as Bcl6 or Pax5 repress the PC phenotype^8,23^, it is then mandatory for B cells to downregulate the expression of the GC profile in order to differentiate in GC-derived PC^11,24^. Once B cells have switched their gene expression profile towards a PC profile, they undergo several rounds of cell division that allow the hypomethylation of the chromatin parts encoding the proteins responsible for the PC profile in an irreversible way^25^. Thus, PC differentiation goes through a remodeling at the epigenetic, transcriptomic and phenotypic profile. These changes allow the functional remodeling of activated and proliferating GC B cells to antibody-secreting PC.

Several studies have tried to identify intermediate states bridging GC B cells to PC *in vivo*. Indeed, cells intermediate in their transcriptomic profile^26^, or LZ GC B cells with PC features such as high expression of *Irf4* in mice^18,27^, have been described in recent years. In humans, transcriptomic studies also highlighted intermediate states expressing PC and cell cycle markers^28–30^. Unfortunately, the scarcity of these states in lymphoid organs have most often precluded their in-depth study and the differentiation trajectory between selected GC B cells and differentiated antibody-producing cells remains uncharted. *In silico* modeling and imaging^31^ and forced selection in *in vivo* models^4^ have suggested that PC exit from the GC occurred through or proximal to the DZ. Several signals appear to be implicated in PC egress from GC (GPR183 signaling, S1P1 and CXCL12 gradients), but no final mechanism has been identified^32,33^. During early T-dependent responses, plasmablasts accumulate at the GC-T zone interface (GTI), but whether this also occurs for GC-derived PC is not known. PC that are found in the lymph node (LN) medulla, further away from the GC DZ, express less proliferation markers (Ki67^+^) than those that are closer to the DZ, indicating a possible spatio-temporal axis of GC to PC differentiation^34^. In addition, it was recently shown that post-selection PC expand clonally without further SHM^6^, but whether this expansion occurs inside or outside of the GC, or where in the draining LN (dLN), is still unknown.

Here, we identified intermediate differentiation states of GC to PC in mouse dLN after model antigen immunization or respiratory virus infection, that we characterized through integrative analyses of phenotypes, transcriptomes, BCR repertoires, and microanatomic localization. We show that GC-derived PC differentiation in dLN implies local proliferation at the interface between the GC DZ and the medulla.

## RESULTS

### Tracking GC B cells and recent GC emigrants after model immunization

We used a GC B cell fate mapping mouse model, *Aicda-Cre-ERT2 x Rosa26-lox-STOP-lox-eYFP*^35^, to track GC B cells and their recent progeny in draining lymph nodes after subcutaneaous immunization. After immunization with the T-dependent model antigen 4-hydroxy-3-nitrophenylacetyl-conjugated keyhole lympet hemocyanin (NP-KLH) in adjuvant, we gavaged the mice once with tamoxifen to induce the eYFP expression in *Aicda* expressing cells, and analyzed cells in draining lymph nodes (dLN) by flow cytometry at day 14 after immunization, at different time points after tamoxifen gavage (**Figure 1a**). In that model, gavaging with tamoxifen after day 6 ensures that the eYFP expression is triggered almost exclusively in GC B cells. Thus, among live activated IgD^neg^ B cells (**Supplementary Figure 1a**), eYFP^pos^ cells 24 hours after tamoxifen treatment consisted in their vast majority of GL7^+^CD38^-^ GC B cells with no detectable eYFP^pos^ CD138^+^ PC, whereas the latter, representing GC-derived PC, were readily identified 72 hours after tamoxifen treatment (**Figure 1b**) and reached a plateau at 96 hours (**Figure 1c**). Analyzing antigen-binding cells among IgD^neg^ B cell subsets in dLN at day 14 after NP-KLH immunization and 4 days after tamoxifen gavage (**Supplementary Figure 1b**), we found that total GC B cells and PC included on average 20% and 30% NP-PE^+^ cells, respectively, while MBC included less than 5% antigen-binding cells (**Figure 1d**). Restricting the analysis to fate-mapped eYFP^+^ cells resulted in a non-significant trend for higher proportions of NP-binding cells in all subsets, in particular for eYFP^+^ MBC where 15% on average were NP-PE^+^ (**Figure 1d**), indicating that B cells exported from the GC from day 10 to day 14 were enriched in cells that could detectably bind antigen. Thus, the *Aicda-Cre-ERT2 x Rosa26-lox-STOP-lox-eYFP* mouse model enabled us to track the output of recent GC emigrants in dLN after subcutaneous model antigen immunization.

**Figure 1:**
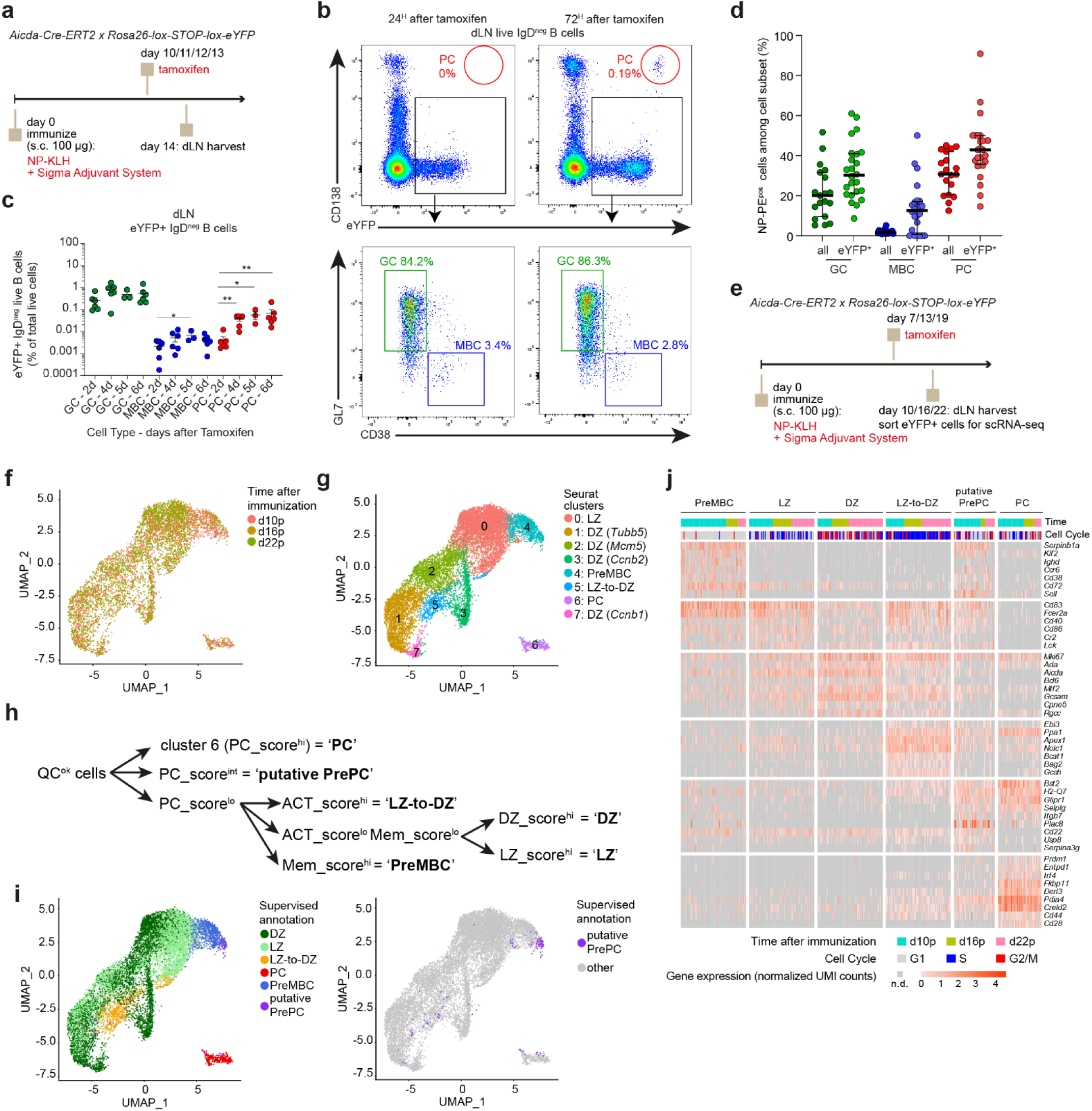
Single-cell RNA-seq analysis of fate-mapped GC B cells and recent GC emigrants identifies putative PrePC in draining LN after immunization. **a**, Experimental design for GC B cell fate mapping. **b**, Flow cytometry gating of eYFP^+^ PC, GC B cells and MBC among IgD^neg^ B cells from dLN in mice 24 hours or 72 hours after fate mapping. Numbers indicate % of parent gate. **c**, Percentage of the indicated eYFP^+^ IgD^neg^ B cell types (GC, MBC or PC) among total live cells in dLN at the indicated times after tamoxifen gavage (2, 4, 5, 6 days). Symbols represent individual mice. * p < 0.05, ** p < 0.01 in Wilcoxon test. **d**, Percentage of antigen-specific cells identified as NP-PE^pos^ by flow cytometry among the indicated cell types. Symbols represent individual mice. Differences between all cells and eYFP^+^ cells are not significant in Dunn’s multiple comparison test after Kruskal Wallis. **e**, Experimental design for scRNA-seq analysis of fate-mapped GC B cells and recent GC emigrants. **f**, UMAP representation of single-cell gene expression profiles of GC and GC-derived cells, colored by time after immunization (d10p: day 10 of primary response). **g**, UMAP representation of single-cell gene expression profiles of GC and GC-derived cells, colored by non-supervised clusters. **h**, Hierarchical gating strategy for gene expression score-based supervised annotation of GC and GC-derived cells (left: all cells; right: highlighting putative PrePC only). **i**, UMAP representation of single-cell gene expression profiles of GC and GC-derived cells, colored by supervised annotation. **j**, Single-cell gene expression heatmap of the indicated marker genes in the different subsets defined by supervised annotation, also indicating the time after primary immunization and cell cycle phase (G1, S, G2/M).

### Single-cell RNA-seq analysis of GC B cells and recent GC emigrants identifies putative PrePC

Using the GC B cell fate mapping mouse model, we generated single-cell RNA-seq (scRNA-seq) gene expression profiles of IgD^neg^ eYFP^+^ B cells from dLN at different time points after primary NP-KLH immunization (**Figure 1e**). After standard quality controls, UMAP embedding of gene expression profiles revealed different clusters of cells contributed from cells samples at all time points (**Figure 1f**). Complementing those scRNA-seq datasets with additional datasets of IgD^neg^ eYFP^+^ B cells from primary or secondary responses to OVA antigen (**Supplementary Figure 1c-d**) or from an independent primary response to NP-KLH antigen (**Supplementary Figure 1d-e**) resulted in similar gene expression profiles. Non-supervised clustering of the single-cell dataset identified 8 clusters that we annotated based on their expressed marker genes (**Figure 1g** and **Supplementary Figure 1g**). Cluster 0 included mostly quiescent cells and corresponded to LZ GC B cells (*Cd83*, *Il4i1*); clusters 1, 2, 3 and 7 expressed distinct combinations of cell cycle associated genes and corresponded to DZ GC B cells; cluster 5 expressed the typical Myc-induced signature of positively selected LZ-to-DZ GC B cells (*Npm1*, *C1qbp*); cluster 4 included mostly quiescent cells and expressed both LZ and MBC markers (*Klf2*, *Serpinb1a*), suggesting those cells corresponded to preMBC or early MBC; and cluster 6 expressed high levels of Ig coding genes and corresponded to PC. Although there was a clear continuum between quiescent LZ GC B cells and preMBC in the low dimensional embedding, the PC cluster was completely separated and we failed to identify transitional states from GC to PC through non-supervised analyses in that dataset.

We reasoned that a supervised approach may be better able to identify rare putative PrePC that would bridge the GC and PC transcriptional states. We thus used a gene signature-based scoring approach to hierarchically “gate” cells in the integrated scRNA-seq dataset (Methods, **Supplementary Table 1**, **Figure 1h** and **Supplementary Figure 1h-i**). In particular, we identified rare cells expressing intermediate levels of a PC-specific gene signature, which we named “putative PrePC”. Other supervised annotations, DZ, LZ, LZ-to-DZ, preMBC and PC were consistent with non-supervised clusters (**Supplementary Figure 1h-i**). PreMBC, LZ, DZ, LZ-to-DZ and PC expressed the expected gene expression programs that have already been described in other mouse GC B cell scRNA-seq datasets^36,37^. Putative PrePC expressed *Bst2*, *H2-Q7*, *Glipr1*, *Selplg*, *Itgb7*, *Plac8*, *Cd22*, *Usp8* and *Serpina3g* among other marker genes (**Figure 1i**). In comparison to recently selected LZ-to-DZ cells, putative PrePC expressed lower levels of *Aicda*, *Bcl6*, *Gcsam* or *Apex1* genes, among others, suggesting that putative PrePC are more advanced in the GC to PC differentiation after positive selection. Interestingly, high expression of *Selplg*, encoding the surface marker PSGL1, suggested that putative PrePC cells may be identified and enriched by flow cytometry.

### Enrichment of putative PrePC by flow cytometry

We thus designed a 14-color flow cytometry panel targeting surface markers and transcription factors characteristic of GC, MBC and PC, and including the PSGL1 marker, which we applied to analyze IgD^neg^ B cells in dLN of *Aicda-Cre-ERT2 x Rosa26-lox-STOP-lox-eYFP* mice previously immunized with NP-KLH and gavaged with tamoxifen (**Figure 2a**). UMAP embedding, based on 10 surface and intracellular markers (Methods) identified clusters of phenotypically defined GC, MBC and PC, as well as cells located in intermediate areas of the low dimensional embedding (**Figure 2b**). We gated cells situated within GC and PC clusters as “GC-to-PC” and inspected their surface phenotype in comparison with the well-defined GC, MBC and PC clusters. GC-to-PC cells expressed high levels of the GC markers GL7, CD19 and B220, high levels of the PC-specific transcription factor IRF4, intermediate levels of the PC marker CD138, and intermediate levels of PSGL1 (**Figure 2c**). Based on the specific surface phenotype of GC-to-PC cells, we thus reverse engineered a gating strategy (**Figure 2d**) that allowed for the FACS-based identification and quantification of putative PrePC cells (**Figure 2e**), which we call “PrePC” for simplicity in the rest of the manuscript. At days 14-16 after primary immunization with NP-KLH, PrePC accounted for approximately 0.08% of IgD^neg^ B cells in dLN (**Figure 2f**). PrePC cells included an average of 18% NP-binding cells, only slightly lower than GC B cells (23%) and PC (24%), and much higher than the proportion of MBC detectably binding NP (2%), suggesting that most PrePC cells were generated in ongoing antigen-specific B cell response (**Figure 2g**). In kinetics experiments with the GC fate mapping model (**Figure 2h**), fate-mapped eYFP^+^ PrePC cells were detected approximately 24 hours earlier than fate-mapped eYFP^+^ PC, as early as 48 hours post-tamoxifen gavage (**Figure 2i**), consistent with their identity of intermediate states in the GC to PC continuum. Although the proportions of PrePC among live IgD^neg^ B cells in dLN were stable for all time points tested after primary immunization (**Supplementary Figure 2a**), we found a significant tendency for increased frequencies of PrePC during secondary responses as compared to primary responses (**Supplementary Figure 2b**).

**Figure 2:**
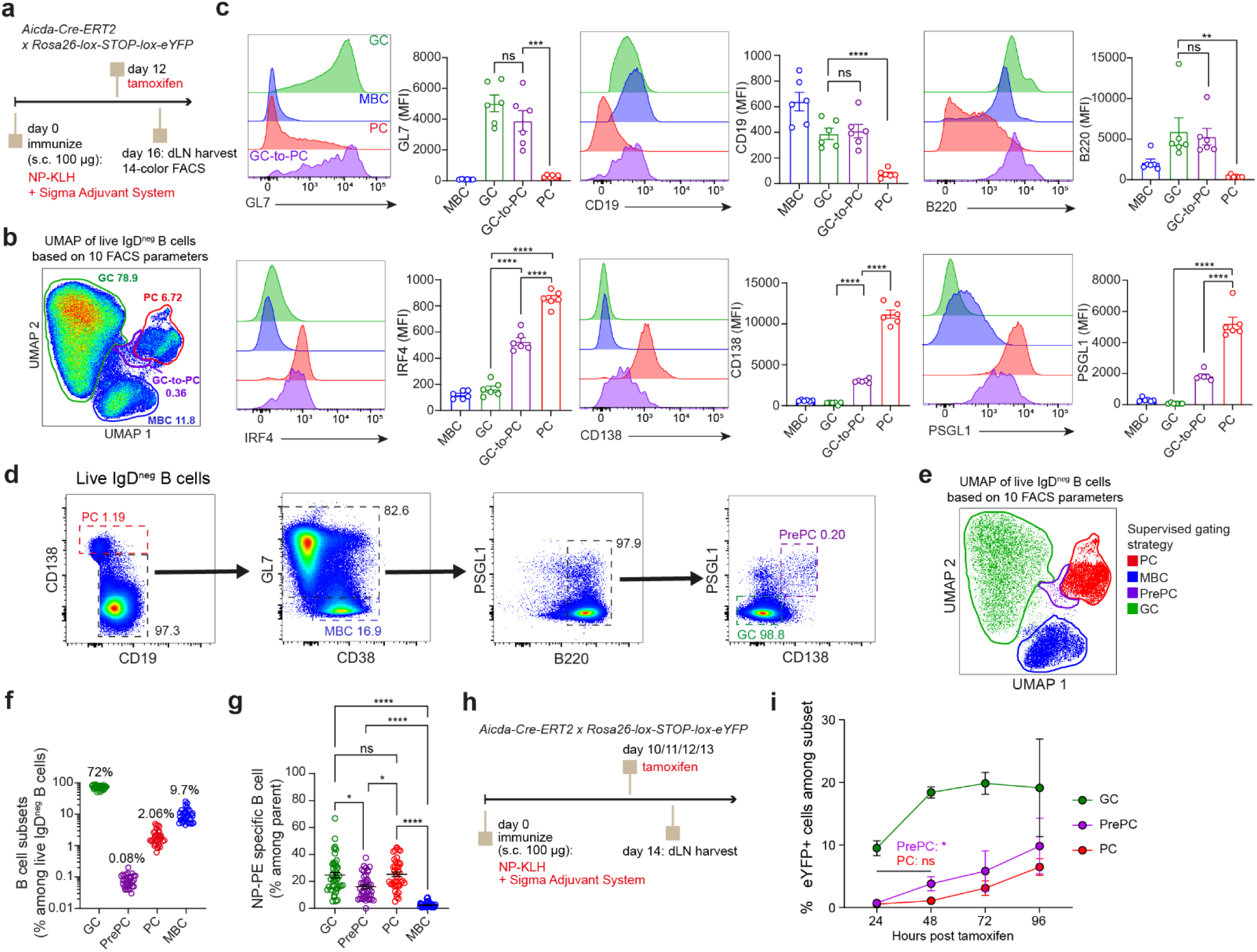
Enrichment of putative PrePC by flow cytometry. **a**, Experimental design for the analysis of putative PrePC phenotype. **b**, UMAP representation of flow cytometry-based protein expression profiles of dLN IgD^neg^ B cells, pseudocolored by embedded cell density. Clusters of GC, PC, MBC and intermediate GC-to-PC cells are circled and annotated. **c**, Mean Fluorescence Intensity (MFI) histogram and quantification for the indicated GC (GL7, CD19, B220; top panels) and PC (IRF4, CD138, PSGL1; bottom panels) protein markers in the indicated cell subsets. ** p<0.01, *** p < 0.001, **** p < 0.0001 in Wilcoxon matched pairs signed rank test. **d**, Gating strategy to enrich for putative PrePC by flow cytometry. **e**, UMAP projection as in b, representing only the cells from the subsets gated in d. **f**, Proportion among dLN IgD^neg^ B cells of cells from the indicated supervised gates defined in d and e. Data compiled from 6 experiments. **g**, Proportion among dLN IgD^neg^ B cells (bottom) of NP-specific cells from the indicated supervised gates defined in d and e. Data compiled from 6 experiments. * p<0.05, **** p < 0.0001, adjusted p-value in Friedman test with Dunn’s multiple comparisons test. **h**, Experimental design for the analysis of fate-mapped GC and GC derived cells at different times after fate mapping. **i**, Percentage of fate-mapped (eYFP+) cells among cells from the indicated phenotypes at different times after tamoxifen-induced fate mapping. Results of statistical testing between percentages at the 24-hour and 48-hour time points for PrePC and PC are displayed on the figure (* p<0.05, ns: not significant, adjusted p-value from Mann-Whitney test after Kruskal-Wallis, with Dunn’s correction).

### Characterization of PrePC cells in the GC to PC differentiation continuum in wild type animals after immunization

In order to gain more insight into the molecular features of PrePC cells, we next used our new gating strategy to enrich and sort phenotypically defined PrePC cells. We first sorted PrePC from wild type animals immunized with NP-KLH and compared them directly to DZ, LZ and PC in FACS-based 5-prime-end single-cell RNA-seq (FB5P-seq)^38^ analysis (**Figure 3a**). After quality control and cell cycle regression of the resulting dataset, low-dimensional UMAP embedding displayed a clear separation between DZ and LZ GC B cells on one side, and PC cells on the other side, with a subset of phenotypically defined PrePC cells bridging the two cell continents (**Figure 3b**). Consistent with our previous analysis on non-enriched cells (**Figure 1i**), single PrePC combined the expression of GC B cell signature genes *Ms4a1*, *Cd19*, *Irf8*, with the expression of positive-selection induced gene *Myc*, and the expression of PC differentiation surface markers and transcription factors *Sdc1*, *Prdm1*, *Irf4*, *Selplg* and *Bst2* (**Figure 3c**). Those cells were also actively proliferating, either in S or G2/M phase of the cell cycle (**Figure 3d**), and expressed high levels of the activation gene expression signature induced upon selection at the LZ-to-DZ state (**Figure 3e**). PC differentiation is characterized by the production of high amounts of antibodies, requiring high level of Ig genes transcription and the induction of a specific endoplasmic reticulum (ER) stress response^33^. PrePC cells expressed intermediate levels of genes involved in the ER stress response (**Figure 3f**), and intermediate levels of Ig transcript counts (**Figure 3g**), when compared with GC B cells and PC. We thus defined a GC to PC continuum of differentiation based on single-cell gene expression of the ER stress module and Ig transcripts counts (**Supplementary Figure 3a-b** and **Figure 3h**), in which most phenotypically defined PrePC bridged the gap between GC B cells and PC. Another feature of PC differentiation, the gradual loss of antigen-presentation capacity on MHC-II^39^, was also intermediate in phenotypically defined PrePC cells (**Supplementary Figure 3c-d**).

**Figure 3:**
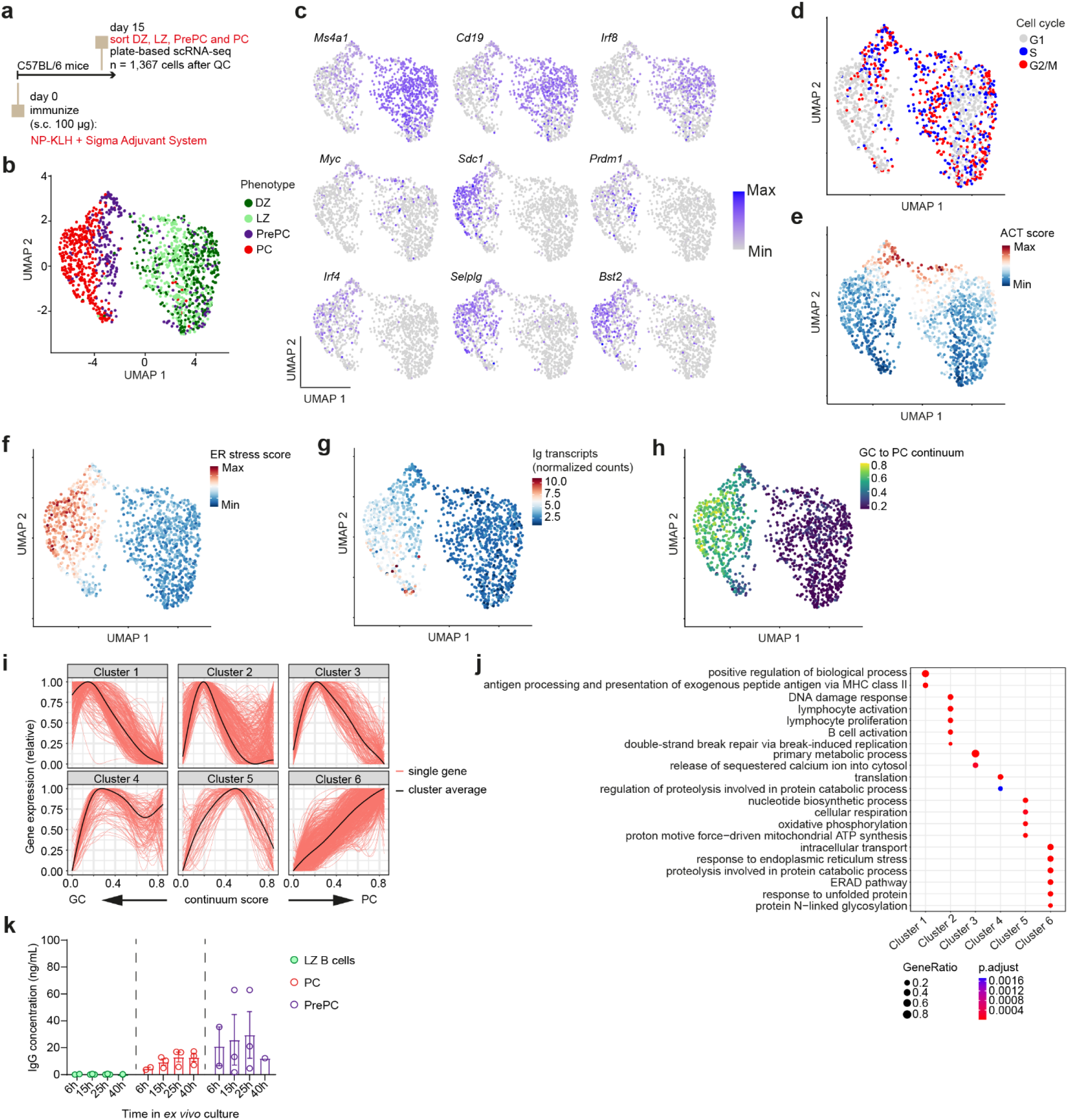
FACS-enriched PrePC cells from wild type immunized animals have an intermediate gene expression profile in the GC-to-PC differentiation continuum. **a**, Experimental design for scRNA-seq of FACS-enriched prePC cells from wild type immunized animals. **b**, UMAP projection of gene expression profiles of FACS-sorted DZ, LZ, PrePC and PC cells, after cell cycle regression. Each dot is a cell, colored by the FACS sorting phenotype. **c**, Feature plots showing the expression of the indicated GC (*Ms4a1*, *Cd19*, *Irf8*, *Myc*) and PC (*Sdc1*, *Prdm1*, *Irf4*, *Selplg*, *Bst2*) marker genes in cells laid out as in b. **d**, UMAP projection as in b, with cells colored by cell cycle status. **e**, UMAP projection as in b, colored by the expression level of the activation gene expression signature. **f**, UMAP projection as in b, colored by the expression level of the ER stress gene expression signature. **g**, UMAP projection as in b, colored by the expression level of Ig transcripts. **h**, UMAP projection as in b, colored by the GC to PC continuum score. **i**, Relative gene expression profile of cells ranked by continuum score, for genes grouped according to their evolution profile (each gene a red line, the average cluster profile a black line). **j**, Results of gene ontology (GO) enrichment analysis for genes in the different evolution clusters defined in i. **k**, IgG concentration in culture supernatant of LZ B cells, PC and PrePC at the indicated times after *ex vivo* cell culture.

Our enriched dataset provided a unique opportunity to map the transcriptional changes that pave the GC to PC differentiation. We thus clustered genes according to the dynamic evolution of their expression as cells progress through the GC to PC continuum (**Figure 3i** and **Supplementary Table 2**) using a method previously developed to analyze the LZ to DZ continuum of GC reactions^40^, and computed the gene ontology enrichment of gene modules that are progressively lost (clusters 1-3) or induced (clusters 4-6) through differentiation (**Figure 3j**). Those analyses revealed that very early metabolic reprogramming (nucleotide biosynthesis process, cellular respiration, oxidative phosphorylation, proton motive force−driven mitochondrial ATP synthesis) preceded the gradual increase in antibody production-associated physiological responses (response to ER stress, response to unfolded protein). Functionally, FACS-sorted PrePC spontaneously secreted detectable amounts of soluble IgG in *ex vivo* cultures (**Figure 3k**). Thus, PrePC cells with the GL7^+^ B220^+^ CD138^int^ PSGL1^+^ phenotype were engaged in the early stages of PC differentiation and had already initiated the antibody production program.

### PrePC phenotype cells from wild type immunized animals contain GC-derived and GC-independent cells

The affinity maturation trajectory of GC B cells and their progeny is in-part traceable through the accumulation of somatic mutations in their Ig gene transcripts^41^. Our PrePC-enriched FB5P-seq analysis retrieved IgH and IgL sequences for the vast majority of single cells (n=965/1367). For better resolution of the different gene expression programs and their associated BCR sequence features, we re-annotated the cell types and states using the supervised annotation method described in **Supplementary Figure 1h**, resulting in the identification of DZ, LZ, LZ-to-DZ, PC, PreMBC and PrePC cells with specific gene expression programs and phenotypes. While close to 90% of DZ, LZ and LZ-to-DZ subsets carried mutations in their IGH and IgL variable genes, only 75% of preMBC and 40% of PrePC and PC were mutated (**Figure 4a**), suggesting that some of those cells were not derived from GC reactions. Unmutated and mutated PrePC and PC were slightly different in isotype usage, with unmutated cells being more frequently IgM-positive: IgM-expressing cells represented approximately 25% of both DZ and LZ cells, less than 5% of positively selected LZ-to-DZ cells, and about 60% of preMBC; in unmutated PC and PrePC, 30% were IgM-positive, as compared to 5% and 10%, respectively, for their mutated counterparts (**Figure 4b**). We identified BCR clonotypes in the dataset and analyzed the compositions in distinct cell types and states of clonal families (groups of 2 or more cells sharing the same IgH and IgL clonotype). We annotated clonal families depending on whether they contained cells from one or several of the cell states defined by our supervised annotation; then we analyzed for each cell state the proportion of cells being allocated to distinct clonal families (**Figure 4c**). About half of DZ and LZ cells were in clonal families comprising either only GC B cells (DZ, LZ and/or LZ-to-DZ), GC and PC, or GC, PC and PrePC cells. By contrast, LZ-to-DZ cells were more frequently clonally associated to GC, PrePC and PC, and PreMBC were most often not clonally related to any other cell in the dataset. About one quarter of PrePC cells were clonally related to both GC and PC cells, but PrePC also included a majority of non-clonal cells. Interestingly, PrePC and PC cells that were clonally related to contemporaneous GC B cells had significantly more somatic hypermutation load than those that were not (**Figure 4d**), confirming that PrePC cells in wild-type animals include both GC-derived and GC-independent cells.

**Figure 4:**
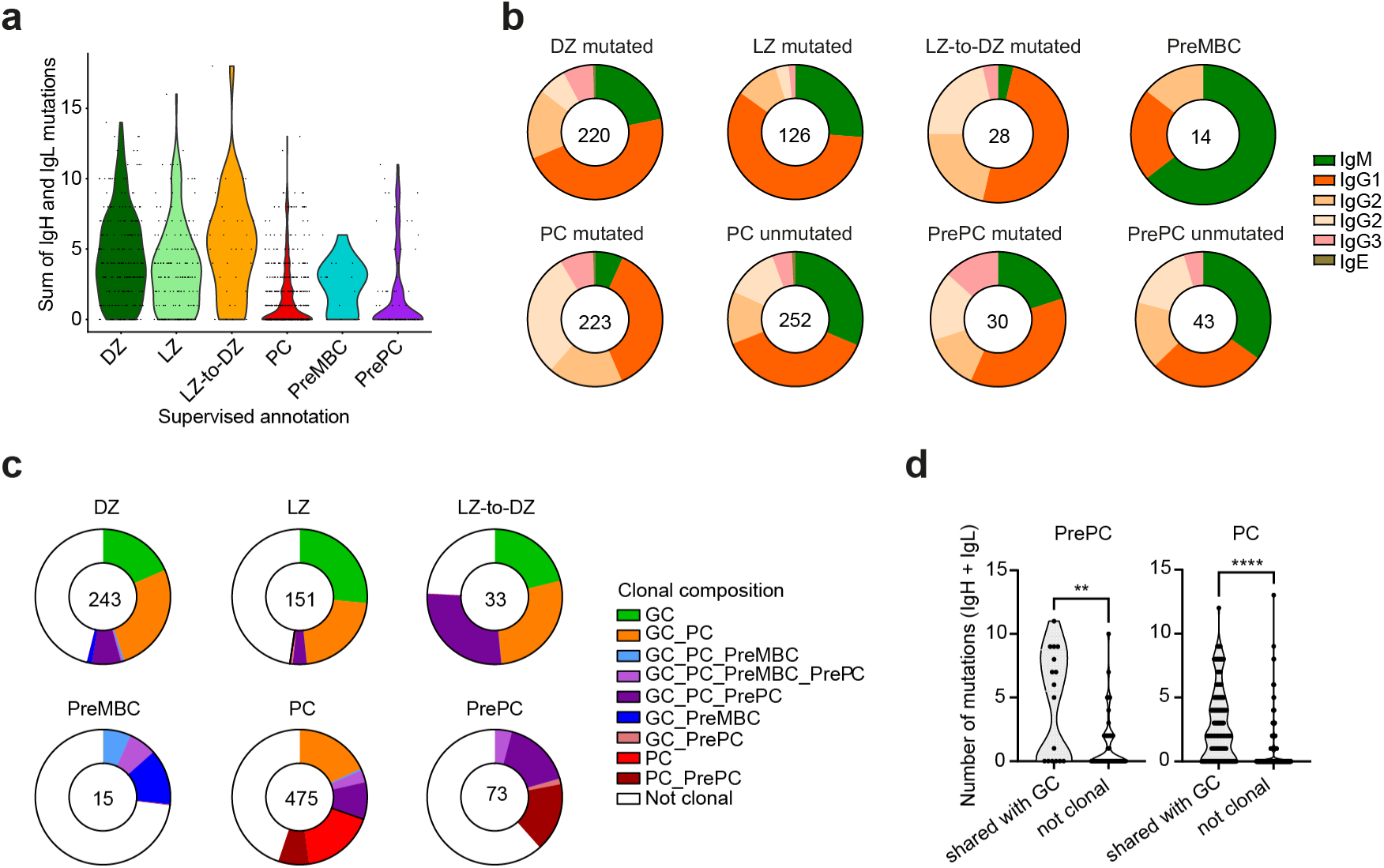
FACS-enriched PrePC cells from wild type immunized animals include both GC-derived and non-GC-derived cells. **a**, Violin plot of BCR mutations (sum of IgH and IgL) in the indicated cell subsets from the dataset described in Figure 3. **b**, Pie charts of the isotype repartition in the indicated cell subsets, with PC and PrePC divided into BCR-unmutated and BCR-mutated. Numbers indicate the number of cells analyzed for each subset. **c**, Pie charts of shared clonotype composition within each subset of the supervised annotation. Numbers indicate the number of cells analyzed for each subset. **d**, Violin plots representing the number of mutations in PrePC (left) and PC (right) from the indicated clonal groups. ** p < 0.01, **** p < 0.0001 in Mann-Whitney non-parametric test.

### Characterization of fate-mapped PrePC cells in the GC to PC differentiation continuum after immunization or infection

In order to focus on GC-derived PrePC and to extend our findings to other types of immune responses, we used the GC fate mapping model to study GC and recent GC emigrant cells, comparing cells from inguinal lymph nodes after subcutaneous NP-KLH immunization with cells from mediastinal lymph nodes after intranasal influenza virus infection (**Figure 5a**). Two weeks after primary activation, infection like protein immunization generated cells with a PrePC phenotype, along with GC, PC and MBC; PrePC and PC were slightly more frequent among activated B cells after infection than after immunization (**Figure 5b**). We sorted fate-mapped live IgD^neg^ eYFP^+^ B cell subsets for plate-based scRNA-seq, enriching for PrePC based on the CD19^+^ GL7^+^ PSGL1^pos^ CD138^lo^ phenotype, as well as DZ (CD19^+^ GL7^+^ CD138^neg^ PSGL1^neg^ CXCR4^hi^ CD86^lo^), LZ (CD19^+^ GL7^+^ PSGL1^neg^ CD138^neg^ CXCR4^lo^ CD86^hi^), PreMBC (CD19^+^ GL7^+^ PSGL1^neg^ CD138^neg^ CD38^+^ CCR6^+^), MBC (CD19^+^ GL7^-^ CD38^+^), PC (CD138^hi^) and CD19^+^ GL7^+^ PSGL1^pos^ CD138^neg^ GC. After quality control of the resulting dataset, low-dimensional UMAP embedding showed that phenotypically defined PrePC from both the NP-KLH and the influenza conditions mapped to a transcriptional space at the interface between GC B cells and PC (**Figure 5c**), also characterized by intermediate levels of GC (CD19, GL7) and PC (CD138, PSGL1) surface proteins (**Figure 5d**). Fate-mapped eYFP^pos^ PrePC and PC expressed BCR with similar levels of somatic hypermutation as contemporaneous DZ and LZ cells (**Figure 5e**) and were biased towards IgG isotypes (**Figure 5f**), consistent with being derived from GC B cells. Thus, the PrePC CD19^+^ GL7^+^ CD138^lo^ PSGL1^+^ phenotype combined with GC fate mapping accurately identified a transitional state in the GC to PC differentiation continuum.

**Figure 5:**
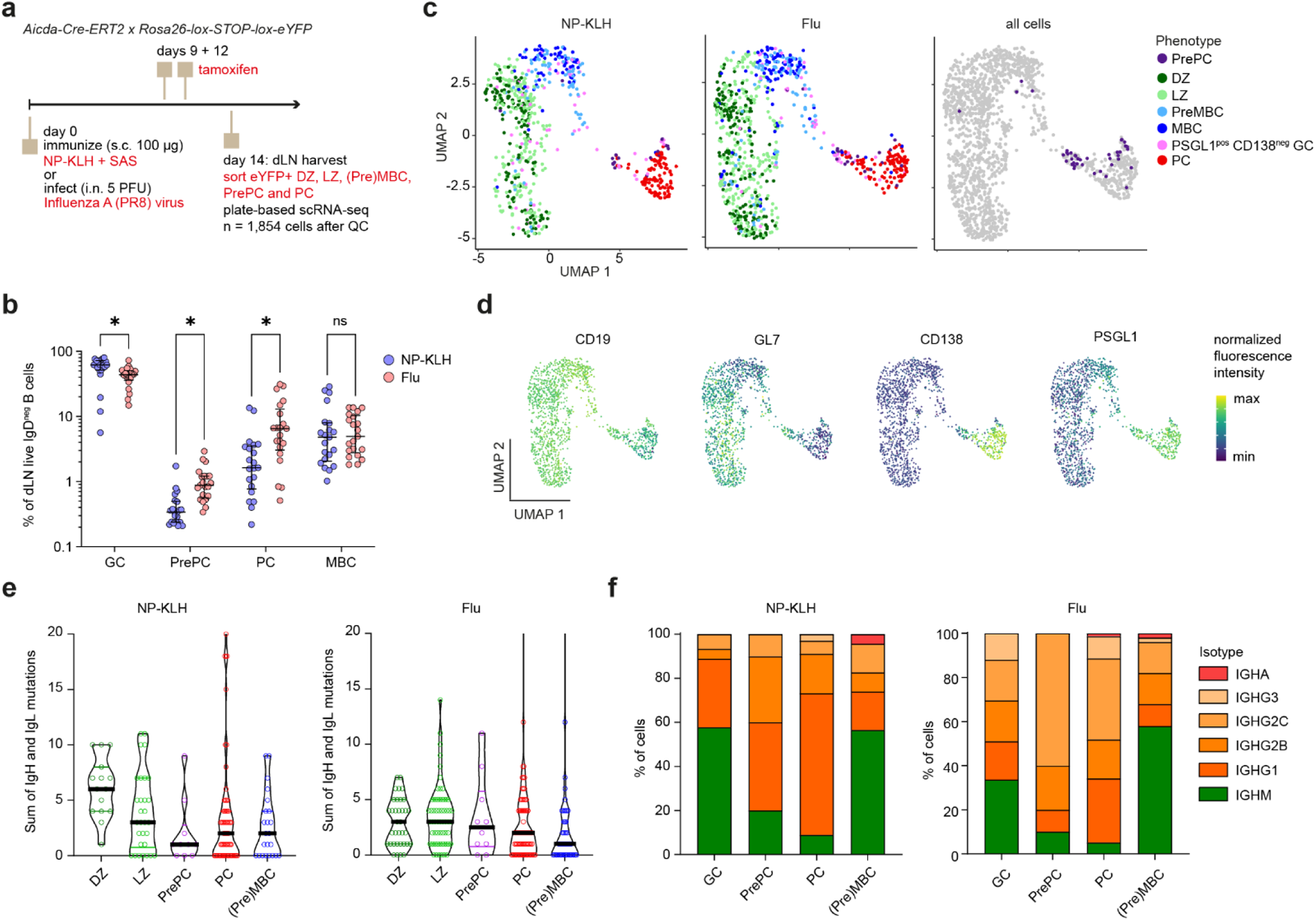
Characterization of fate-mapped FACS-enriched PrePC cells from draining lymph nodes after subcutaneous protein immunization or respiratory influenza virus infection. **a**, Experimental design for scRNA-seq analysis of fate-mapped GC and GC-derived cells after NP-KLH immunization or influenza virus infection. **b**, Percentage of the indicated cell types (GC, PrePC, PC or MBC) among live IgD^neg^ B cells in dLN 14 days after NP-KLH immunization (blue symbols) or Influenza virus infection (red symbols). Each symbol represents a mouse. * p<0.05 adjusted p-value from Wilcoxon matched-pairs signed rank test. **c**, UMAP projection of gene expression profiles of FACS-sorted fate-mapped PrePC, DZ, LZ, PreMBC, PSGL1^pos^ CD138^neg^ GC B cells and PC cells from NP-KLH immunized (left) or influenza virus infected (middle) animals. Each dot is a cell, colored by the FACS sorting phenotype. Right panel highlights PrePC cells from both activation conditions. **d**, Feature plots showing the expression of the indicated surface proteins in cells laid out as in c, based on index sorting analysis. **e**, Violin plot of BCR mutations (sum of IgH and IgL) in the indicated cell subsets from NP-KLH immunized (left) or influenza virus infected (right) animals. The (Pre)MBC annotation combines PreMBC and MBC phenotypes. **f**, Bar plots of the isotype repartition for the indicated cell subsets from NP-KLH immunized (left) or influenza virus infected (right) animals. The (Pre)MBC annotation combines PreMBC and MBC phenotypes.

### Recent GC-derived PC differentiate and proliferate at the GC DZ-medulla interface

While GC B cells migrate within the confined GC micro-environment of draining lymph nodes, GC-derived MBC and PC are physically excluded from GC structures^42,43^. In order to infer whether the early stages of PC differentiation occur within or outside the GC micro-environment, we first analyzed the expression levels of genes expressing key homing receptors and adhesion proteins in our fate-mapped scRNA-seq dataset (**Figure 6a**). Cells in the PrePC cluster expressed a unique combination of those genes (*Cxcr5*^neg^ *S1pr2*^neg^ *Ccr7*^neg^ *Sell*^low^ *S1pr1*^low^ *Selplg*^hi^ *Itga4*^pos^ *Itgb1*^pos^ *Cxcr3*^pos^) which was distinct from that expressed by LZ, DZ, (Pre)MBC or PC, suggesting that PrePC cells may be localized in a specific compartment in the dLN outside of GC (*S1pr2*^neg^) and not in the B follicle (*Cxcr5*^neg^ *Sell*^low^) or T cell zone (*Ccr7*^neg^).

**Figure 6:**
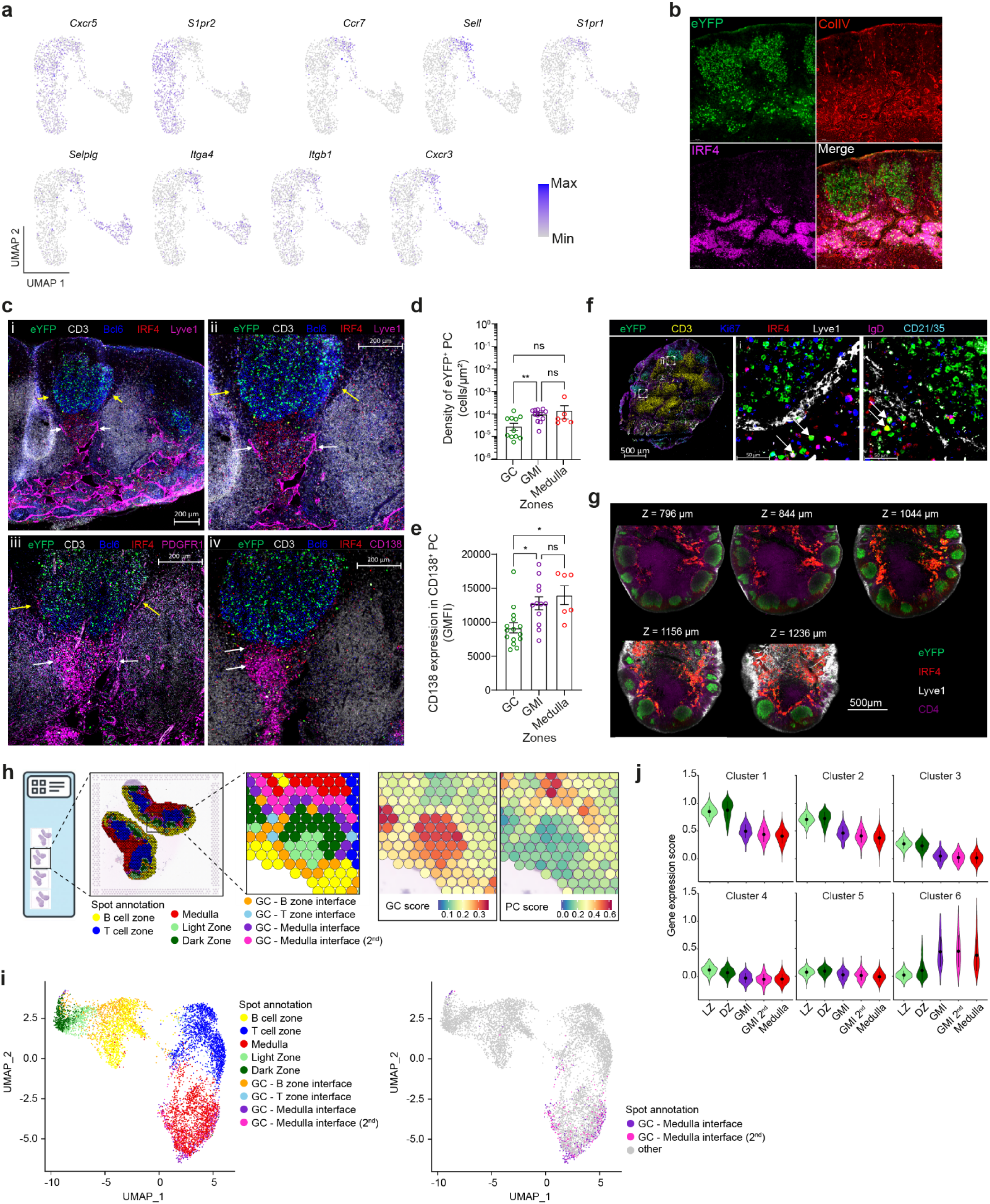
Recent GC-derived PC accumulate at the GC-medulla interface. **a**, Feature plots showing the expression of the indicated homing markers in cells laid out as in Figure 4c. **b**, Confocal microscopy image of the indicated markers in a dLN section (14 days after NP-KLH immunization, 4 days after tamoxifen), focusing on two GC and the DZ-proximal area. **c**, Spectral confocal microscopy images of serial dLN sections (14 days after primary NP-KLH immunization, 3 days after tamoxifen) stained for the indicated markers. Images show a broad view around a GC (i) and a zoomed in view highlighting the GTI and the GMI (yellow and white arrows in (ii) and (iii), respectively) as well as the distinct populations of CD138^lo^ and CD138^+^ cells (white arrows in (iv)). **d**, Density of GC-derived PC (IRF4^+^ cells expressing the GC fate-mapping eYFP) quantified from spectral confocal microscopy images after segmentation in the indicated zones. Each dot represents a zone, combining analyses from 2 dLN in 2 distinct experiments. **e**, Expression level of CD138 (geometric mean fluorescence intensity) in mature PC (IRF4^+^ CD138^+^ cells) in the indicated zones. Each dot represents a zone, combining analyses from 2 dLN in 2 distinct experiments. **f**, Spectral confocal microscopy images of a representative dLN from an influenza virus-infected mouse (14 days after intranasal infection, 4 days after tamoxifen) stained for the indicated markers, highlighting the GMI from two GCs in (i) and (ii). Zoomed-in views of (i) and (ii) only show the eYFP, Ki67, IRF4 and Lyve1 markers. White arrows point to recent GC-derived PC (IRF4^+^ eYFP^+^ cells). **g**, Lightsheet microscopy images of a whole dLN stained with the indicated markers, extracted from different Z-positions as indicated above images. **h**, Experimental design of the spot-based spatial transcriptomics analysis of dLN, and semi-supervised annotation of spots based on unsupervised spot clustering, gene expression scores, and neighboring spot identity. **i**, Gene expression-based UMAP projection of spatial transcriptomics spots, colored by spot annotation. Bottom view highlights only spots annotated as GMI and GMI 2^nd^. **j**, Violin plots of gene expression scores in spots from the indicated zones for the gene clusters identified based on their evolution on the GC to PC continuum in Figure 3i.

To assess the location of GC-derived PC differentiation, we used high-resolution microscopy to precisely map the location of eYFP^+^ IRF4^+^ cells, which we defined as recently differentiated PC, in dLN two weeks after immunization and 4 days after tamoxifen fate mapping. We found clusters of eYFP^+^ IRF4^+^ cells accumulating in collagen-IV-rich areas resembling PC-rich medulla in close proximity to the GC DZ (**Figure 6b**). Those areas, where eYFP^+^ IRF4^+^ cells were surrounded by Lyve1^+^ lymphatic endothelial cells and PDGFR1^+^ stromal cells (**Figure 6c**), connected the GC DZ to the medulla and were thus termed GC-medulla interface (GMI). The density of eYFP^+^ IRF4^+^ cells in the GMI and medulla was similar (**Figure 6d**). In some tissue sections, the eYFP^+^ IRF4^+^ cells in close proximity to the GC DZ expressed lower levels of CD138 than those more distal in the medullary areas (**Figure 6c**, panel *iv*), suggesting less advanced differentiation; that observation was not statistically significant when considering all GCs from multiple dLN sections (**Figure 6e**). We detected similar DZ-proximal Lyve1^+^ GMI environments accumulating fate-mapped eYFP^+^ IRF4^+^ in mediastinal lymph nodes from influenza infected mice (**Figure 6f**). Since we did not find a GMI area surrounding every GC in every tissue section, we reasoned that the medulla may connect with the GC DZ only at certain contact areas that are not always captured in a 10-20 µm tissue section. Using whole-LN 3D-imaging with light sheet microscopy we could indeed determine that every GC in a dLN included a GMI contact area, but that the area was not found at all depths (**Figure 6g** and **Supplementary Movie**).

To characterize whether the GMI is the site of early PC differentiation and maturation, we performed whole transcriptome spot-based spatial transcriptomics (10x Genomics Visium) analysis of 4 sections of the same 2 dLN, separated by approximately 50µm in depth (**Figure 6h**). After quality control, we annotated spots with a semi-supervised approach: first using non-supervised spot clustering and marker genes to identify spots corresponding to B follicles, T zones, medulla, GC LZ and GC DZ; then defining three zones in close contact to the GC (Methods), the GC-T zone interface (GTI), the GC-B zone interface (GBI), and the GC-medulla interface (GMI); and labeling medulla spots in proximity to GMI as 2^nd^ order GMI (**Figure 6h-i**). GMI expressed a gene expression profile that was distinct from GC-distal medulla (**Figure 6i**), with 35 differentially expressed genes upregulated in GMI *vs.* medulla, of which 28 genes were also upregulated in DZ *vs.* medulla (including cell cycle related genes *Tuba1b*, *Hmces*, *Mcm5*) and 7 were not related to the DZ environment (*Psap*, *Mzb1*, *Ighm*, *Ctsb*, *Emc4*, *Clu*, *Exosc5*) (**Supplementary Table 3**). The identity of those genes suggested that the gene expression changes characterizing GC to PC differentiation was occurring along a LZ to DZ to GMI to GMI 2^nd^ to medulla spatial axis. Indeed, clusters of genes characterizing the continuum of GC to PC differentiation (**Figure 3i**) followed spatially ordered expression gradients consistent with our hypothesis (**Figure 6j**). In particular, genes from clusters 1 and 2 characteristic of GC B cells somatic hypermutation and proliferation (**Figure 3j**) peaked in DZ and decreased gradually from DZ to GMI to GMI 2^nd^ to medulla; by contrast, genes from cluster 6 characteristic of PC differentiation (**Figure 3j**) were highly expressed from the GMI, increasing further in GMI 2^nd^, and slightly decreasing in medulla areas.

Our scRNA-seq data and recently published observations^6,44^ indicated that early differentiating PrePC, or recently differentiated PC proliferate extensively. In confocal images, a majority of eYFP^+^ IRF4^+^ at the GMI expressed Ki67, a hallmark of active proliferation (**Figure 7a**). Consistently, cell cycle related gene expression was higher in GMI than GMI 2^nd^ or medulla areas in our spatial transcriptomics dataset (**Figure 7b**). Proliferating fate-mapped eYFP^+^ IRF4^+^ cells appeared at the GMI from 48 hours after tamoxifen-induced fate mapping and could only be detected later at low density in the more distal medulla (**Figure 7c-f**). MacLean et al. have shown that GC-derived PC proliferation is regulated by the affinity of their BCR for antigen^44^. In our model, fate-mapped eYFP^pos^ PrePC and eYFP^pos^ PC that detectably bound the NP antigen tended to include more Ki67+ proliferating cells, although the differences did not reach statistical significance (**Figure 7g-h**). This suggests that affinity-regulated positive selection controls the proliferation of post-GC PC from the PrePC intermediate stage.

**Figure 7:**
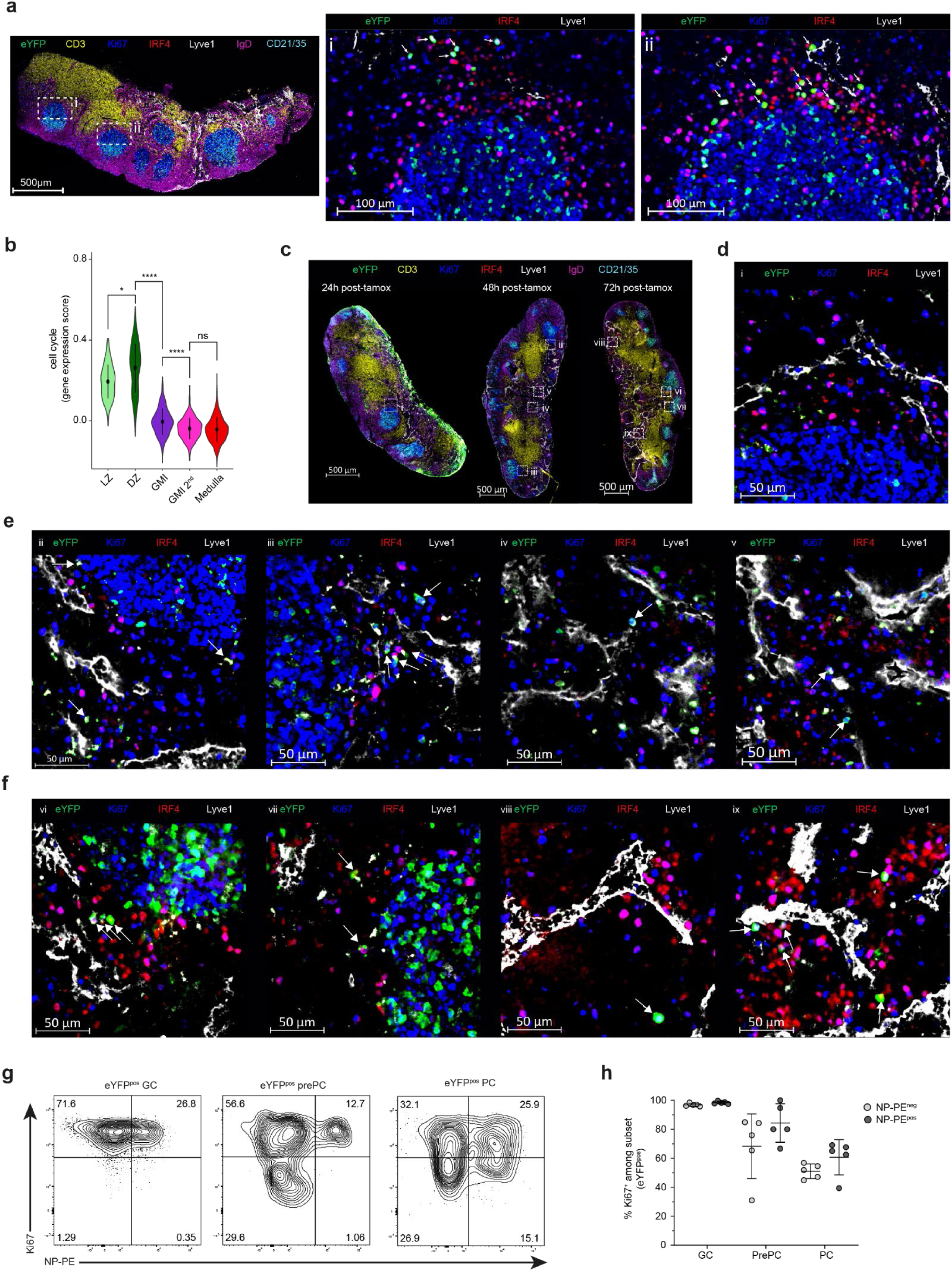
Recent GC-derived PC proliferate at the GC-medulla interface. **a**, Spectral confocal microscopy images of a dLN section (14 days after primary NP-KLH immunization, 3 days after tamoxifen) stained for the indicated markers. Images show the whole dLN section (left) and zoomed-in DZ proximal areas of two GCs (GC i: middle; GC ii: right). Arrows in the zoomed-in panels indicate triple positive eYFP^+^ IRF4^+^ Ki67^+^ GC-derived proliferating PC. **b**, Violin plot of cell cycle gene expression scores in spots from the indicated zones defined in Figure 6h-i. * p<0.05, **** p<0.0001, adjusted p-value from Mann-Whitney test with Dunn’s correction. **c**, Spectral confocal microscopy images of dLN sections (14 days after primary NP-KLH immunization) at the indicated times after tamoxifen induction of GC fate mapping, stained for the indicated markers. White squares with roman numerals indicate GMI and medulla areas shown with higher magnification in subsequent panels. **d**, Spectral confocal microscopy images of a GMI (area i, 14 days after primary NP-KLH immunization) 24 hours after tamoxifen induction of GC fate mapping, stained for the indicated markers. **e**, Spectral confocal microscopy images of 2 GMI areas (ii and iii) and 2 medulla areas (iv and v), 14 days after primary NP-KLH immunization and 48 hours after tamoxifen induction of GC fate mapping. **f,** Spectral confocal microscopy image of 2 GMI areas (vi and vii) and 2 medulla areas (viii and ix), 14 days after primary NP-KLH immunization and 72 hours after tamoxifen induction of GC fate mapping. **g,** Representative flow cytometry contour plots showing NP-PE binding (x-axis) and Ki67 expression (y-axis) in the indicated eYFP^pos^ cell subsets from dLN 14 days after NP-KLH immunization (4 days after tamoxifen). Numbers in quadrants indicate percentages. **h,** Percentage of Ki67^+^ cells among the indicated eYFP^pos^ cell subsets, comparing NP-PE^neg^ (gray dots) and NP-PE^pos^ (black dots) cells.

Overall, we have characterized the GMI, a specific dLN microenvironment defined as an extension of the medulla contacting the GC DZ at some level, as the main environment for exit and proliferation of recent GC-derived PC.

## DISCUSSION

Our study provides a fine characterization of rare PrePC intermediate states in the GC to PC differentiation trajectory in draining LN after immunization or infection. We showed that PrePC cells expressed a mixed phenotype expressing both GC and early PC markers in phenotypic and transcriptomic analyses; they were also actively cycling, expressed intermediate levels of MHC-II genes, had initiated metabolic reprogramming and started upregulating UPR programs, and produced antibodies *ex vivo* without any additional signal. Those data indicate that we have identified late intermediate states in the GC to PC differentiation trajectory. PrePC expressed a distinctive repertoire of homing markers, suggesting they had already homed or were homing to extra-GC areas in draining LN. We found recently GC-exported PC proliferating just outside of the GC DZ, at the interface with medullary tissue microenvironments that connected every GC to the deep medulla. The GC-medulla interface contained Lyve1^+^ and PDGFR1^+^ stromal cells that may serve as a maturation niche for cells in the GC to PC differentiation trajectory.

Several studies have identified PrePC states as LZ GC B cells expressing high levels of *Irf4* in mice^45,46^, and cells expressing both PC and cell cycle markers in humans^7,12,47^. Those studies lacked other information such as their surface phenotype, function, and *in situ* localization, making it difficult to compare them directly with our definition of PrePC. Recent studies in human tonsils has identified pre-plasmablasts that appeared integrated in the differentiation trajectory from GC to PC^28,29,48^. Indeed, the authors found cells that were intermediate in their transcriptomic profile, and shared clones with both GC B cells and PC. Even though the authors did not focus their analyses on these states, we could identify some common features with the PrePC we have identified in our study in mice (*e.g.* expression of PC lineage genes like *IRF4*, *PRDM1*, *XBP1*, *FKBP11*, and cell cycle progression), highlighting some evolutionary conserved modules activated at the GC-to-PC transition. In another high resolution human tonsil atlas^49^, GC-derived PC branched both from LZ GCB cells and activated LZ-to-DZ B cells, the latter representing the likely counterparts of the activated PrePC we describe here in the mouse LN. Another study characterized human PrePC in an *ex vivo* MBC differentiation model using a combination of scRNAseq and ATACseq^50^. Although they originated from blood MBC, and not LN GC B cells, PrePC in that study were highly heterogenous and integrated in a continuum of differentiation with stepwise induction of the unfolded protein response, and with AP-1 family transcription factors like BATF playing a likely role at the onset of differentiation. In our study, PrePC branched from the LZ-to-DZ GC B cell state, where the transcription factor BATF has been shown to play a key role for metabolic refuelling induced by T cell help^51^. Given that the PrePC cells that we identify with our gating strategy have already initiated transcriptional rewiring linked to antibody production and are already producing soluble antibodies, whereas LZ-to-DZ cells do not, it is likely that our strategy is missing earlier intermediates in the GC to PrePC differentiation. Although the IgD^neg^ CD19^+^ GL7^+^ PSGL1^+^ CD138^neg^ phenotype seemed like an interesting approach for enriching such early intermediates, our scRNA-seq analyses in Figure 4c show that those cells are heterogeneous transcriptionally and not a sufficiently robust option. Altogether, our current results on PrePC are consistent with previous studies, refine the molecular, phenotypic and BCR repertoire characterization of those rare intermediate states, and introduce a gating strategy for enriching those cells from mouse lymph node by flow cytometry, which should be an asset for future studies.

*In silico* modeling experiments have suggested that PC exit the GC from the DZ^31^. Several homing receptors appear to be implicated in PC egress from GC (Gpr183, S1PR1, CXCR4), but the precise route that PrePC or PC use to exit GCs remained unidentified^32,33^. Based on gene expression, we found that PrePC were likely in the process of egress from the GC, but their scarcity precluded their robust identification by confocal microscopy in LN tissue sections. Instead, we analyzed recent GC-derived PC that were eYFP^+^ IRF4^+^ 3-4 days after tamoxifen-induced fate mapping of GC B cells. GC-derived PC accumulated at the border of the GC. In previous studies, PC differentiated after early GC-independent B cell activation accumulated at the interface between the nascent GC and the T zone^45,52^. In our study, we showed that GC-derived PC were in fact accumulating and proliferating in DZ-proximal microenvironments characterized by a gene expression and cellular composition similar to the LN medulla. Notably, the GC-medulla interface was surrounded by Lyve1^+^ cells resembling medullary sinus LEC^53^, and contained numerous PDGFR1^+^ stromal cells with high gene expression scores for previously described medullary stromal cell subsets (Nr4a1^+^ and Inmt^+^ stromal cells, data not shown)^52^. Medullary fibroblastic reticular cells have been shown to provide essential survival and maturation cues to plasma cells in lymph nodes^54^, but the fact that those cells may be in such a close proximity to the GC DZ has remained unnoticed in imaging studies focusing on B follicle stromal remodeling^55,56^. We attribute that to the fact that most 2D images of thin tissue sections failed to capture the GMI, which bridged medulla and GC DZ only at certain z-depth for a given GC structure. We did not detect GMI areas in non-immunized lymph nodes (unpublished observation), and other studies have shown a tight separation between medulla and B cell follicles in naïve LN^54,57^. However, adjuvanted immunization or infection induces lymphangiogenesis and stromal remodeling^54,58^ close to activated B cell follicles, which may initiate the spontaneous organization of a GMI area that supports early PC proliferation, maturation and migration.

In our experiments, close to 25% of LZ-to-DZ cells were clonally related to contemporaneous PC or PrePC, compared to less than 10% for LZ or DZ GC B cells (**Figure 4c**), suggesting that PrePC derive more closely from GC B cells clones selected at the LZ-to-DZ stage. Thus, our study implies that, after being selected in the LZ, a fraction of LZ-to-DZ state GC B cells commit to the PrePC state, proliferate at the external border of the DZ and mature locally at the GC-medulla interface. For selected GC B cells recycling into the DZ compartment, it has been shown that higher affinity induced more cycles of proliferation in recycled DZ B cells^59,60^. Similarly, high affinity and increased signals from the BCR and T_FH_ help induces robust proliferation of GC-derived PC^44^. Our analyses suggest that this process of affinity-regulated post-selection PrePC maturation and proliferation occurs in specific niches where interactions with specific components of the GMI micro-environment likely plays a key role in the process, thereby regulating overall serological antibody affinity maturation.

## EXPERIMENTAL METHODS

### Mouse models

*Aicda-Cre-ERT2 x Rosa26-lox-STOP-lox-eYFP* mice^35^ were bred at the Centre d’Immuno-Phenomique, (Marseille, France), and transferred to the animal care facility of Centre d’Immunologie de Marseille-Luminy for experiments. C57BL/6 mice were purchased from Janvier Labs (Le Genest-Saint-Isle, France). All mice were maintained in the CIML mouse facility under specific pathogen-free conditions. Experimental procedures were conducted in agreement with French and European guidelines for animal care under the authorization number APAFIS #30945-2021040807508680 (NP-KLH immunization) and APAFIS #48826-2024040816466800 (Influenza virus infection), following review and approval by the local animal ethics committee in Marseille. Mice were used regardless of sex, at ages greater than 7 weeks and less than 3 months.

Mice were immunized with either 100µg of NP-KLH at 1µg/µL emulsified with Sigma Adjuvant System (SAS) at a 1:1 (v:v) ratio, or 100µg chicken ovalbumin (OVA) at 1µg/µl emulsified with Alum at a 1:1 (v:v) ratio, subcutaneously at the base of the tail, 50µl on each side. Mice were anesthetized i.p. with Ketamine/Xylazine (100 mg/kg body) and intranasally infected with 5 PFU of Influenza virus A/Puerto Rico/8/1934 (PR8) H1N1 strain as described^61^. For induction of the Cre-ERT2-mediated labelling, we gavaged the mice once or twice, as indicated in experimental diagrams and figure legends, with 5mg of tamoxifen (TS648-1G, Sigma) in 200µl of peanut oil (P2144-250 ML, Sigma) for NP-KLH immunizations or 100 µl of corn oil (405435000, Thermo Scientific Chemicals), at least 6 days after immunization. Mice were euthanized between 10 days and 28 days post-immunization (prime or boost) according to the experiments.

### Flow cytometry

Single-cell suspensions from draining lymph nodes were washed and resuspended in FACS buffer (5% fetal calf serum, 2mM EDTA, 5% Brilliant Stain Buffer Plus in PBS 1X) at a concentration of 100 million of cells per ml. Cells were first incubated with FcBlock (Biolegend) for 10 min on ice. Then, cells were incubated with a mix of antibodies (see table1 below) conjugated with fluorochromes 30 min on ice. Cells were washed in PBS, and incubated with the Live/Dead Fixable Aqua Dead Cell Stain (Thermofisher) for 10 min on ice. For intranuclear stainings, cells were washed and permeabilized using the FoxP3 permeabilization kit (eBioscience) during 30min, then washed again in the permeabilization buffer and incubated with intracellular antibodies for 45min at RT. Cells were finally washed in permeabilization buffer and resuspended in FACS buffer. Cell suspensions were analyzed on the LSR II (with UV laser) or LSR Fortessa X20 cytometers (Becton Dickinson).

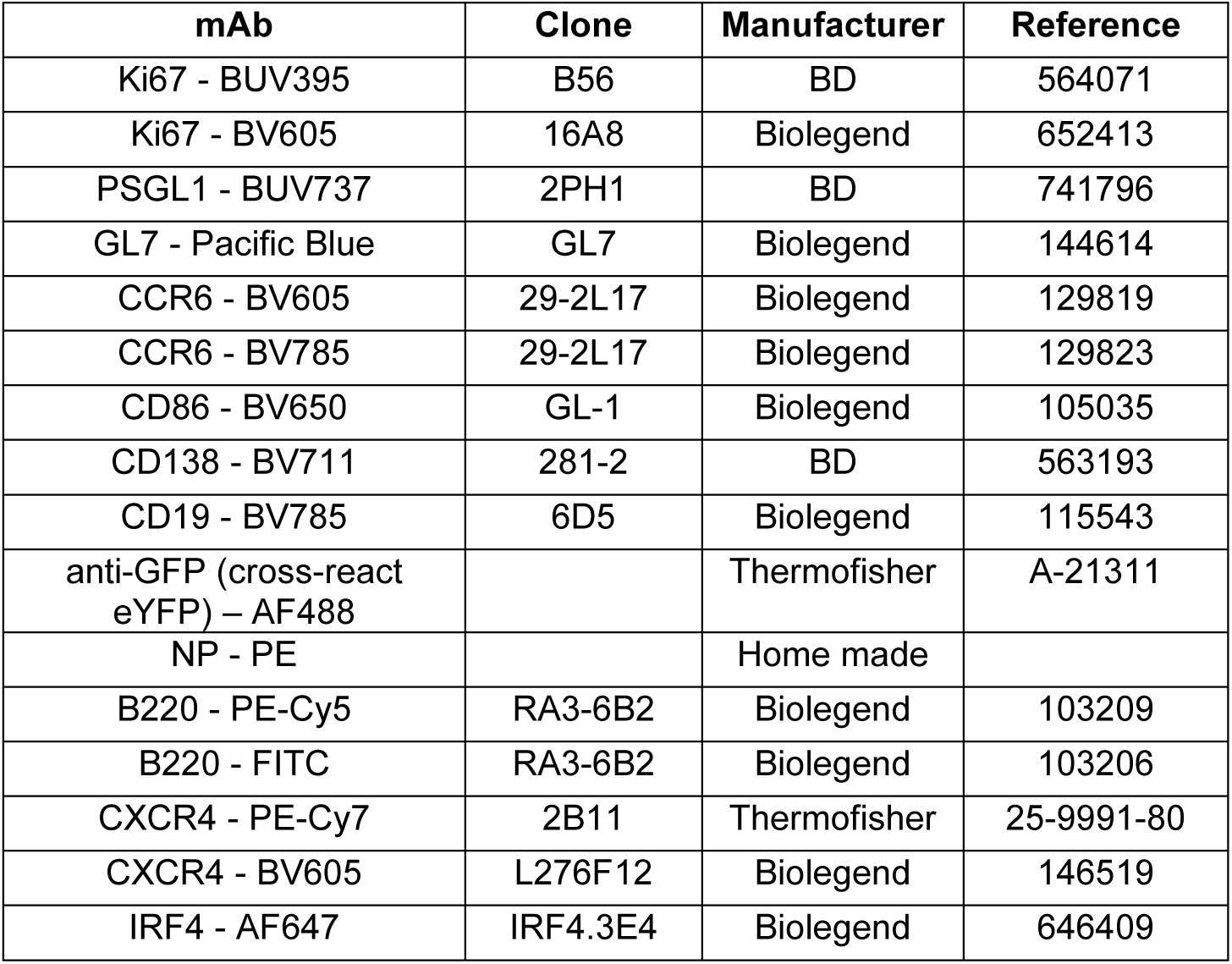
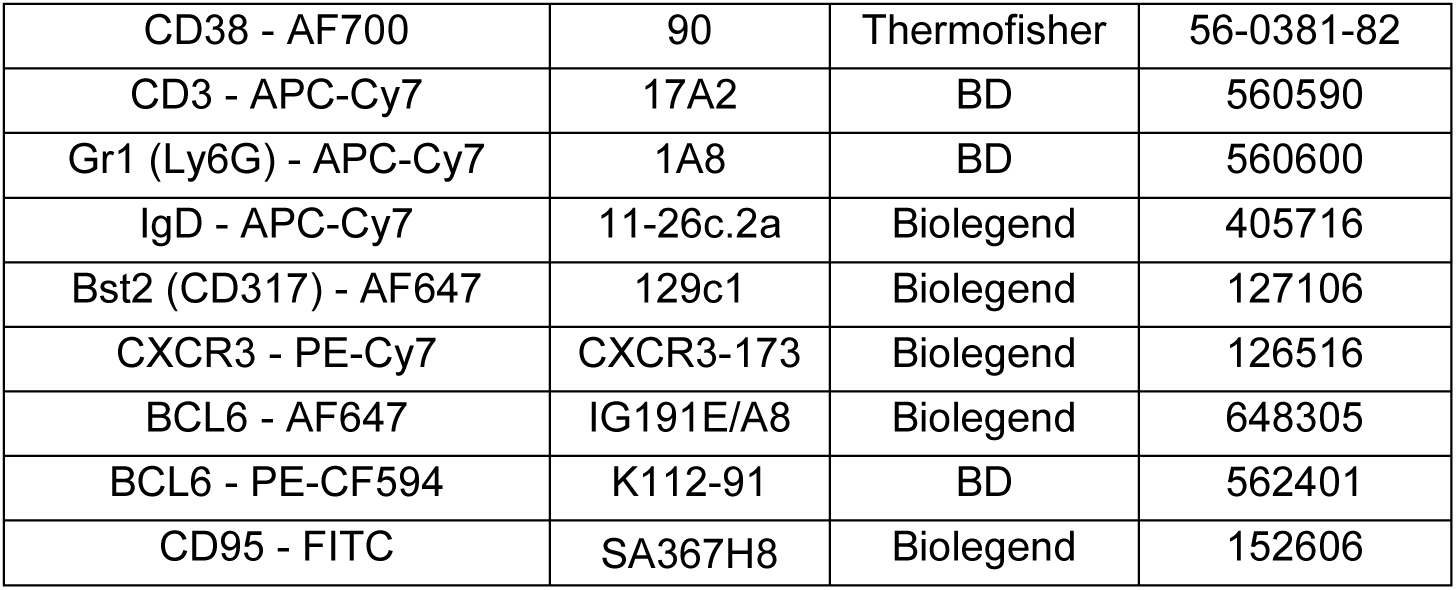

| mAb | Clone | Manufacturer | Reference |
| --- | --- | --- | --- |
| Ki67 - BUV395 | B56 | BD | 564071 |
| Ki67 - BV605 | 16A8 | Biolegend | 652413 |
| PSGL1 - BUV737 | 2PH1 | BD | 741796 |
| GL7 - Pacific Blue | GL7 | Biolegend | 144614 |
| CCR6 - BV605 | 29-2L17 | Biolegend | 129819 |
| CCR6 - BV785 | 29-2L17 | Biolegend | 129823 |
| CD86 - BV650 | GL-1 | Biolegend | 105035 |
| CD138 - BV711 | 281-2 | BD | 563193 |
| CD19 - BV785 | 6D5 | Biolegend | 115543 |
| anti-GFP (cross-react eYFP) – AF488 |  | Thermofisher | A-21311 |
| NP - PE |  | Home made |  |
| B220 - PE-Cy5 | RA3-6B2 | Biolegend | 103209 |
| B220 - FITC | RA3-6B2 | Biolegend | 103206 |
| CXCR4 - PE-Cy7 | 2B11 | Thermofisher | 25-9991-80 |
| CXCR4 - BV605 | L276F12 | Biolegend | 146519 |
| IRF4 - AF647 | IRF4.3E4 | Biolegend | 646409 |
| CD38 - AF700 | 90 | Thermofisher | 56-0381-82 |
| CD3 - APC-Cy7 | 17A2 | BD | 560590 |
| Gr1 (Ly6G) - APC-Cy7 | 1A8 | BD | 560600 |
| IgD - APC-Cy7 | 11-26c.2a | Biolegend | 405716 |
| Bst2 (CD317) - AF647 | 129c1 | Biolegend | 127106 |
| CXCR3 - PE-Cy7 | CXCR3-173 | Biolegend | 126516 |
| BCL6 - AF647 | IG191E/A8 | Biolegend | 648305 |
| BCL6 - PE-CF594 | K112-91 | BD | 562401 |
| CD95 - FITC | SA367H8 | Biolegend | 152606 |

For single-cell sorting, cells were pre-enriched using the “Pan B Cell Isolation Kit II mouse” enrichment kit from Miltenyi Biotec (ref. 130-104-443) in which we added a biotinylated anti-IgD antibody for further enrichment of IgD^neg^ B cells (Biolegend ref. 405734). Cells were prepared as mentioned in the protocol provided and passed through LS columns according to the manufacturer’s instructions. We collected the negative fraction and processed the cells according to the classical extracellular staining protocol, using antibodies described in the table below. Cells were sorted on the BD Influx (FB5P-seq experiments, see below) or the BD FACS Aria III SORP (Flash-FB5P-seq experiments, see below) cell sorter, in 96-well PCR plates, with index-sorting mode for recording the fluorescence parameters associated to each sorted cell.

The panel used for the FB5P-seq experiments was the following:

| mAb | Clone | Manufacturer | Reference |
| --- | --- | --- | --- |
| B220 - FITC | RA3-6B2 | Biolegend | 103205 |
| CXCR4 - PerCP-eF710 | 2B11 | Thermofisher | 46-9991-82 |
| GL7 - Pacific Blue | GL7 | Biolegend | 144614 |
| PSGL1 (CD162) - AF647 | 2PH1 | BD | 562806 |
| CD3 - APC-Cy7 | 17A2 | BD | 560590 |
| Gr1 (Ly6G) - APC-Cy7 | 1A8 | BD | 560600 |
| IgD - APC-Cy7 | 11-26c.2a | Biolegend | 405716 |
| CD38 - PE | 90 | Invitrogen | 12-0381-82 |
| CD19 - PE-Dazzle594 | 6D5 | Biolegend | 115554 |
| CD86 - PE-Cy7 | GL-1 | Thermofisher | 12-0862-82 |

The panel used for the Flash-FB5P-seq experiments was the following:

| mAb | Clone | Manufacturer | Reference |
| --- | --- | --- | --- |
| PSGL1 - BVU737 | 2PH1 | BD | 562806 |
| GL7 - Pacific Blue | GL7 | Biolegend | 144614 |
| CXCR4 - BV605 | L276F12 | Biolegend | 146519 |
| CD86 - BV650 | GL-1 | Biolegend | 105035 |
| CD138 - BV711 | 281-2 | BD | 563193 |
| CCR6 - BV785 | 29-2L17 | Biolegend | 129823 |
| CD19 - PE-Dazzle594 | 6D5 | Biolegend | 115554 |
| CXCR3 - PE-Cy7 | CXCR3-173 | Biolegend | 126516 |
| CD38 - AF700 | 90 | Thermofisher | 56-0381-82 |
| CD3 - APC-Cy7 | 17A2 | BD | 560590 |
| Gr1 (Ly6G) - APC-Cy7 | 1A8 | BD | 560600 |
| IgD - APC-Cy7 | 11-26c.2a | Biolegend | 405716 |
| NP - PE (NP-KLH) |  | Home made |  |
| HA – PE (Flu) |  | Home made |  |
| NP - APC (Flu) |  | Home made |  |

Data were analysed using FlowJo (v10.8.1 and v10.10.0).

### FB5P-seq

For the experiments described in Supplementary Figure 1c, Figure 3 and Supplementary Figure 3, we used the FB5P-seq protocol as previously described by Attaf et al.^38^. Individual cells were sorted into a 96-well PCR plate, with each well containing 2 µL of lysis buffer. Index sort mode was activated to record the fluorescence intensities of all markers for each individual cell. Flow cytometry standard (FCS) files from the index sort were analyzed using FlowJo software, and compensation parameters were exported as CSV tables for subsequent bioinformatic analysis. Immediately after sorting, plates containing individual cells were stored at -80°C until further processing. Following thawing, reverse transcription was performed, and the resulting cDNA was preamplified for 22 cycles. Libraries were then prepared according to the FB5P-seq protocol. The FB5P-seq data were processed to generate both a single-cell gene count matrix and single-cell B cell receptor (BCR) repertoire sequences for B cell analysis. Two separate bioinformatic pipelines were employed for gene expression and repertoire analysis, as detailed in Attaf et al.^38^.

### Flash-FB5P-seq

For the experiments described in Figure 5, we used an updated version of the FB5P-seq protocol, termed Flash-FB5P-seq, as described in Cardon et al.^62^. This new protocol implemented advances developed by Hahaut et al. in the Flash-seq protocol^63^. Cells were sorted in 2 µl lysis mix composed of 0.1% Triton X-100 (0.02 µl), 1.2 U/µl recombinant RNAse inhibitor (0.06 µl), 6 mM dNTP mix (0.48 µl), 1.6 mM RT primer (5′-TGCGGTATCTAAAGCGGTGAGTTTTTTTTTTTTTTTTTTTTTTTTTTTTTT*V*N-3′) (0.36 µl), 0.0125 pg/µl ERCC spike-in mix (0.05 µl), 9 mM dCTP (0.18 µl), 1 M betaine (0.4 µl), 1.2 mM DTT (0.024 µl) and PCR-grade H_2_O (0.426 µl). Plates were conserved at -70 °C until sorting, thawed at RT for cell sorting, and immediately refrozen on dry ice then conserved at -70 °C until further processing. Reverse transcription with template switching and cDNA amplification were performed in a single step. Each plate containing cells in lysis mix was thawed on ice, heated at 72 °C for 3 min then immediately placed on ice. To each well containing 2 µl cell lysate, we added 8 µl of an RT-PCR mix composed of 2 U/µl Superscript IV reverse transcriptase (0.1 µl), 0.8 U/µl recombinant RNAse inhibitor (0.2 µl), 1X KAPA HiFi ReadyMix (5 µl), 4.8 mM DTT (0.48 µl), 0.8 M betaine (1.6 µl), 9.2 mM MgCl_2_ (0.092 µl), 0.1 µM forward PCR primer (5’-AGACGTGTGCTCTTCCGAT*C*T-3′) (0.1 µl), 0.1 µM reverse PCR primer (5′-TGCGGTATCTAAAGCGGTG*A*G-3′) (0.1 µl), 1.84 µM biotinylated barcoding template switching oligonucleotide (5′-biotin-AGACGTGTGCTCTTCCGATCTXXXXXXXXNNNNNNNNCAGCArGrGrG-3′, where XXXXXXXX are well-specific barcodes as published^38^, NNNNNNNN are random nucleotides serving as Unique Molecular Identifiers, and rG are riboguanosine RNA bases) (0.184 µl) and PCR-grade H_2_O (0.144 µl). Then plates were incubated in a thermal cycler 60 min at 50 °C, 3 min at 98 °C, and 22 cycles of 20 s at 98 °C, 20 s at 67 °C and 6 min at 72 °C. We then pooled 5 µl amplified cDNA from all wells into one tube per plate and performed 0.6X SPRI bead-based purification. Sequencing libraries were prepared from 800 pg cDNA per plate with the Illumina Nextera XT protocol, modified to target sequencing reads to the 5’ end of cDNA, as described^38^. Before sequencing, pooled indexed libraries were depleted from ribosomal RNA-derived cDNA molecules by using the SEQoia RiboDepletion Kit (Biorad) according to the manufacturer’s instructions. Flash-FB5P-seq libraries were sequenced on an Illumina NextSeq2000 using P3 100 cycle kit and targeting 5×10^5^-1×10^6^ reads per cell in paired-end single-index mode with the following configuration: Read1 (gene template) 114 cycles, Read i7 (plate barcode) 8 cycles, Read2 (cell barcode and Unique Molecular Identifier) 16 cycles.

### 10x 5’ scRNA-seq

For experiments described in Figure 1e-f and Supplementary Figure 1e-f, cells from draining lymph nodes were washed and resuspended in FACS buffer (PBS containing 5% FCS and 2 mM EDTA) at a concentration of 10^8^ cells/ml. Samples from different mice and different time points were processed separately using cell hashing as described^64^. Cells were individually stained with distinct barcoded anti-mouse CD45 antibodies (in-house conjugated) in FACS buffer for 30 minutes on ice. Subsequently, cells were washed and stained with a mix of primary antibodies, then Live/Dead Fixable Aqua Dead Cell Stain (Thermofisher). Live cells of interest (IgD^neg^ eYFP^+^ PC and B cells) from each sample were bulk-sorted using a BD Influx cell sorter. PC and non-PC cells were captured in distinct wells for droplet-based scRNA-seq to maximize the recovery of BCR sequence information from non-PC cells. Within each fraction, cells from different samples were pooled and loaded for the 10x Genomics Single Cell 5’ v2 workflow. Libraries were prepared according to the manufacturer’s instructions with modifications for generating B cell receptor (BCR) sequencing libraries. Following cDNA amplification, SPRI select beads were used to separate the large cDNA fraction derived from cellular mRNAs (retained on beads) from the hashtag oligonucleotide (HTO)-containing fraction (in supernatant). For the mRNA-derived cDNA fraction, 50 ng were used to generate the transcriptome library, and 10-20 ng were used for BCR library construction. Gene expression libraries were prepared according to the manufacturer’s instructions. For BCR libraries, heavy and light chain cDNAs were amplified by two rounds of PCR (6 cycles + 8 cycles) using external primers recommended by 10x Genomics. Approximately 800 pg of purified amplified cDNA was tagmented using the Nextera XT DNA Sample Preparation kit (Illumina) and amplified for 12 cycles using the SI-PCR forward primer (10x Genomics) and a Nextera i7 reverse primer (Illumina). For the HTO-containing fraction, 5 ng were used to generate the HTO library. The resulting libraries were pooled and sequenced on an Illumina NextSeq2000 platform with single-indexed paired-end kits following the manufacturer’s guidelines.

### Ex vivo culture and ELISA

Single-cell suspensions FACS-sorted from draining lymph nodes were washed and resuspended in culture medium (10% FCS, 0.1% 2-Mercapto-ethanol, 1% non-essential amino acids, 1% Sodium Pyruvate, 1% HEPES buffer; 1% L-Glutamine, 1% Peniciline-Streptomycin in RPMI 1640) and placed in culture for different times at 37°C, 5% CO_2_. Cell culture supernatants were collected and stored at -80°C until analysis. Mouse IgG concentrations were determined using the LSBio Mouse IgG ELISA Kit (Catalog No. LS-F10451) according to the manufacturer’s instructions.

### Spatial transcriptomics

Freshly dissected draining lymph nodes were dryed on absorbent paper, embedded in OCT, snap frozen in isopentane over dry ice, and stored at -80°C until processing. On the day of the experiment, 10 µm-thick cryosections were prepared from the region of interest using a cryostat. Four sections separated by 50µm in depth were placed on a Visium slide within the designated capture areas. Hematoxylin and eosin (H&E) staining was performed according to the manufacturer’s guidelines with the following modifications: hematoxylin incubation for 30 seconds, bluing buffer for 5 seconds, and eosin incubation for 1 minute 30 seconds, all at room temperature. Following washing, brightfield images of the stained sections were acquired before proceeding to subsequent steps of the 10x Genomics Visium Spatial Gene Expression protocol. The tissue was embedded in OCT medium and a 10 µm section was cut on a cryostat and deposited on the capture area of the Visium slide following the guidelines of the 10x Genomics Visium Spatial Gene Expression protocol. Briefly, we performed permeabilization (18 min), reverse transcription, second strand synthesis, denaturation, cDNA amplification (16 cycles of PCR), and library construction according to the manufacturer’s instructions. The resulting libraries were sequenced on an Illumina NextSeq2000, generating an average of 120 million reads per library (Read 1: 28 cycles, Read i7: 10 cycles, Read i5: 10 cycles, Read 2: 79 cycles).

### Confocal microscopy

Draining lymph nodes were harvested from immunized mice and fixed in antigen fix solution (DiaPath, ref. P0014) for 2 hours at 4°C. Samples were subsequently washed in phosphate-buffered saline (PBS) and cryoprotected in 30% sucrose overnight. The tissue was then embedded in optimal cutting temperature (OCT) compound and snap-frozen in isopentane. Cryosections (20 µm thickness) were prepared using a cryostat and stored at -20°C until staining. For immunofluorescence staining, sections were rehydrated in 1X PBS for 10 minutes. Non-specific binding was blocked by incubating sections for 30 minutes at room temperature in a blocking solution containing 0.1% Triton X-100, 1% fetal calf serum (FCS), 1% bovine serum albumin (BSA), and 1% serum from the host species of the secondary antibody in 1X PBS. Primary antibodies (see table below) were diluted in blocking solution, and sections were incubated overnight at 4°C. Following washing, slides were mounted using ProLong Gold antifade reagent (Invitrogen, ref. P36930). Confocal and spectral images were acquired using a Zeiss LSM 980 confocal microscope. Image processing for conventional analysis was performed using Zen software, while quantitative analysis was conducted using *QuPath* software (see "*Quantitative analysis of confocal microscopy images*" section for details).

| mAb | Clone | Manufacturer | Reference |
| --- | --- | --- | --- |
| Lyve1 - eF570 | ALY7 | Thermofisher | 41-0443-82 |
| Ki67 - eF660 | SolA15 | Thermofisher | 50-5698-82 |
| IRF4 - AF594 | IRF4.3E4 | Biolegend | 646409 |
| CD3 - eF450 | 17A2 | Thermofisher | 48-0032-82 |
| CD21/35 - AF700 | 7E9 | Biolegend | 123431 |
| IgD - SNIR 685 | 11-26c.2a | Biolegend | 405749 |
| PDGFR1 - PE | 18A2 | Biolegend | 323605 |
| anti-GFP (cross-react<br>eYFP) - AF488 |  | Thermofisher | A-21311 |
| Bcl6 - AF647 | IG191E/A8 | Biolegend | 648306 |
| CD138 - PE | 281-2 | Biolegend | 142504 |
| Rabbit anti-mouse Col IV |  | Abcam | Ab19808 |
| Donkey anti-rabbit IgG -<br>Cy3 |  | Jackson<br>ImmunoResearch | 711-165-152 |

### Lightsheet microscopy

Draining lymph nodes were harvested and fixed overnight in 0.4% paraformaldehyde (Electron Microscopy Science, ref. 15714) diluted in phosphate-buffered saline (PBS). Immunostaining was performed using the Hubrecht protocol^65^. Organs were washed in PBS and incubated overnight at 4°C in blocking buffer (PBS containing 5% donkey serum, 1% rehydrated milk, and 0.4% Triton X-100). Samples were then incubated for 5 days with primary antibodies (see table below) diluted in PBS containing 1% rehydrated milk, 0.4% Triton X-100, 3% donkey serum, and 3% mouse serum. Following incubation, samples were washed in PBS with 0.4% Triton X-100 for 1 day. Secondary antibody incubation was performed similarly for 5 days, followed by another 1-day wash in PBS. Samples were embedded in 1% low-melting agarose and subsequently dehydrated through a graded methanol series (20%, 40%, 60%, 80%, and 2× 100% in PBS) for 1 hour per concentration at room temperature. Lymph nodes were then incubated overnight in 100% dehydrated methanol at room temperature. Clearing was initiated by incubating samples in a 1:1 mixture of methanol and BABB (benzyl alcohol and benzyl benzoate at a 1:2 ratio, Sigma ref. 305197 and Fisher Scientific ref. 10654752), followed by overnight incubation in pure BABB. Finally, samples were transferred to ethyl cinnamate for storage until imaging. Imaging was performed using a LaVision Ultramicroscope II (Miltenyi Biotec). Image stacks were acquired with a step size of 6 µm at x2 magnification using an optic zoom with a numerical aperture of 9 µm. Three-dimensional reconstruction and analysis of image stacks were conducted using IMARIS software (Version 9.1.0, Bitplane).

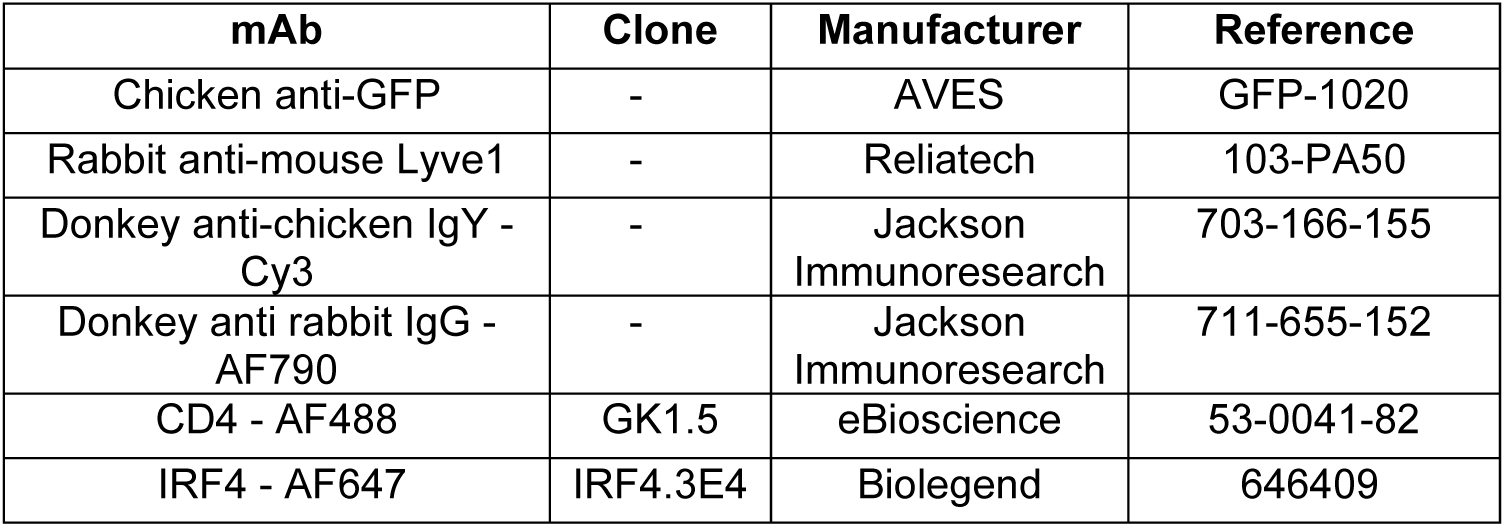

## COMPUTATIONAL METHODS

### Flow cytometry data analysis

The FlowJo UMAP extension was downloaded from FlowJo Exchange Website and used according to the default parameters (euclidean distance, nearest neighbors = 15, minimum distance = 0.5, number of components = 2). The UMAP presented in Figure 2b was computed based on the following compensated parameters: IRF4, Bst2, CD38, GL7, CD138, CD19, PSGL1, Bcl6, B220, CCR6.

### Pre-processing of scRNA-seq datasets

#### 10x Genomics 5’-end sequencing

Raw fastq files from gene expression libraries were processed using *Cell Ranger* and *Cell Ranger VDJ* (v3.0.1 for dataset #2, v6.1.2 for dataset #3), with alignment on the mm10 and vdj_GRCm38_alts_ensembl-7.0.0 reference genomes, respectively. Quality control was performed on each dataset independently to remove poor quality cells based on UMI counts, number of genes detected, percentage of transcripts from mitochondrial genes, and percentage of transcripts from ribosomal protein coding genes. For each cell, gene expression UMI count values were log-normalized with Seurat *NormalizeData* with a scale factor of 10,000 to generate normalized UMI count matrices. HTO barcodes for sample demultiplexing after hashing were counted using CITE-seq-count and were normalized for each cell using a centered log ratio (CLR) transformation across cells implemented in the Seurat function *NormalizeData*. Hashtags were demultiplexed using *MULTIseqDemux* function and barcodes assigned as doublets or negative were excluded from further analysis.

#### FB5P-seq

We used a custom bioinformatics pipeline to process fastq files and generate single-cell gene expression matrices and BCR sequence files as previously described^38^. Detailed instructions for running the FB5P-seq bioinformatics pipeline can be found at https://github.com/MilpiedLab/FB5P-seq. Quality control was performed on each dataset independently to remove poor quality cells based on UMI counts, number of genes detected, ERCC spike-in quantification accuracy, and percentage of transcripts from mitochondrial genes. For each cell, gene expression UMI count values were log-normalized with Seurat *NormalizeData* with a scale factor of 10,000 to generate normalized UMI count matrices.

Index-sorting FCS files were visualized in FlowJo software and compensated parameters values were exported in CSV tables for further processing. For visualization on linear scales in the R programming software, we applied the hyperbolic arcsine transformation on fluorescence parameters^66^.

For BCR sequence reconstruction, the outputs of the FB5P-seq pipeline were further processed and filtered with custom R scripts. For each cell, reconstructed contigs corresponding to the same V(D)J rearrangement were merged, keeping the largest sequence for further analysis. We discarded contigs with no constant region identified in-frame with the V(D)J rearrangement. In cases where several contigs corresponding to the same BCR chain had passed the above filters, we retained the contig with the highest expression level. BCR metadata from the *MigMap* and *Blastn* annotations were appended to the gene expression and index sorting metadata for each cell.

#### Flash-FB5P-seq

We then used a custom bioinformatics pipeline to process fastq files and generate single-cell gene expression matrices and BCR sequence files. Briefly, reads were processed with the zUMIs pipeline^67^ to generate the gene expression UMI count matrix, and with the TRUST4 pipeline^68^ to reconstruct BCR sequences. Quality control was performed as described for FB5P-seq data (above) to remove poor quality cells. The BCR data output from the TRUST4 pipeline was analyzed with IMGT HighV-QUEST to quantify somatic hypermutation.

### Non-supervised analysis

For the analysis described in Figure 1f-g and Supplementary Figure 1d-g, the datasets after quality control and normalization were integrated in *Seurat* using *SelectIntegrationFeatures* (2000 features, excluding BCR coding genes), *FindIntegrationAchors*, and *IntegrateData* (30 components). After integration, we used *RunPCA* (30 components), *FindNeighbors* (30 components) and *FindClusters* (resolution 0.3) for non-supervised clustering, and *RunUMAP* (30 components) for visualization. Marker genes were computed with *FindAllMarkers* (Wilcoxon assay) and top markers visualized as a *Seurat* dot plot. For the analyses described in Figure 3 and Figure 5, we used a standard *Seurat* analysis workflow, including cell cycle regression for the FB5P-seq dataset.

### Supervised annotation

Supervised annotation of scRNA-seq datasets were performed as described in Figure 1h and Supplementary Figure 1h. We used the *AddModuleScore* function to compute gene expression scores for every cell in the datasets for the indicated signatures (Supplementary Table 1). Thresholds for “gating” (Supplementary Figure 1h) were defined empirically to optimize concordance between supervised annotation and non-supervised clustering. The continuum score in Figure 3h was computed by fitting a linear regression on the distribution of B cells in the scatter plot of ER stress score (x-axis) and Normalized Ig transcripts UMI counts (y-axis), and projecting data points on the regression line (Supplementary Figure 3b).

### Gene ontology

The evolution profile of genes along the differentiation continuum was computed by applying a kernel smoother on the distribution of data, for each gene expressed in more than 25% cells in the dataset (n=1834 genes after excluding mitochondrial and ribosomal genes). The derivatives of the resulting profiles were hierarchically clustered (pearson’s correlation distance) to define groups of co-evolving genes in the continuum. Lists of genes grouped in distinct evolution clusters were then submitted to gene ontology analysis using *gprofiler2*, with default settings, and computing only the results for the “BP: biological processes” categories. The results of multiple gene clusters were appended in a single plot using *ggplot2*.

### BCR-seq analysis

Single-cell BCR-seq data were further analyzed with Change-O^69^ to compute clonotypes. The intersection between cell clonotype identities and supervised annotations based on gene expression was used to compute the clonal compositions displayed in Figure 4c.

### Spatial Transcriptomics analysis

Spatial transcriptomics FASTQ files were aligned to the mm10 reference genome using *SpaceRanger* v1.3.1. Downstream analysis was performed using the *Seurat* R package v4.2.1, employing log-normalization and the Louvain clustering algorithm. Annotation of the four merged slices was conducted iteratively, beginning with a clustering resolution of 0.1 (50 principal components) to identify main areas (B cell zone, T cell zone, and medulla), followed by a resolution of 0.2 (50 principal components) to reveal germinal centers. GC spots were further subclustered for light zone (LZ) and dark zone (DZ) segregation (resolution 0.2, 30 principal components). Border regions were annotated based on neighboring spots, defining GC-T zone interface (GTI), GC-medulla interface (GMI), and GC-B zone interface (GBI) based on neighbor spots annotation. The second order GC-medulla interface (GMI 2^nd^) was defined as Medulla spots with a first order GMI neighbor spot. Annotations were projected onto a UMAP calculated using 50 principal components from the complete dataset. Gene signatures were scored with Seurat *AddModuleScore*.

### Quantitative analysis of confocal microscopy images

For cell detection and quantification in confocal microscopy images (Figure 6d-e), we used the *QuPath* analysis platform. We annotated manually the GC and medulla areas based on the eYFP staining for the GC and on the IRF4 staining for the medulla. We defined the GC-medulla interface (GMI) as a 50µm-wide border surrounding the GC and intersecting the medulla annotation. We imported *Cellpose2coloc* in *QuPath* to segment the cells based on the different channels selected, which generated different cell segmentation masks. We determined the positivity thresholds for defining cell subsets, and used them in the colocalization analysis using a custom script. We exported the measurements and performed quantification analyses in GraphPad Prism. All scripts are publicly available here: https://github.com/Imagimm-CIML/Detection-of-B-cell-subtypes-in-a-draining-lymph-node-using-a-mask-colocalization-approach?tab=readme-ov-file#readme.

### Statistical analyses

Statistical analyses were performed using Graphpad Prism or R softwares with tests and p-value significance criteria detailed in the figure legends.

## ACKNOWLEDGEMENTS

We are grateful to all past and present members of the “Integrative B cell Immunology” lab at Centre d’Immunologie de Marseille-Luminy (CIML) for useful discussions, to Marc Bajénoff for stimulating discussions on lymph node stromal cells, and to Hugues Lelouard for his expertise on spectral confocal microscopy. We acknowledge the Computational Biology, Biostatistics and Modeling (CB2M) hub at CIML for helpful discussions and comments. We thank the Flow Cytometry Core Facility, and the Imagimm facility of CIML. We thank the animal care facility of CIML. We acknowledge Centre de Calcul Intensif d’Aix-Marseille for granting access to its high-performance computing resources. This work was supported by grants from ANR (ANR-17-CE15-0009-01 “MoDEx-GC” and ANR-23-CE15-0025-01 “GCselection”) to P.M. This work was supported by institutional grants from INSERM, CNRS and Aix-Marseille University to the CIML. L.B., L.Br. and N.A. were supported by fellowships from the French Ministry of Research and Higher Education.

## AUTHORSHIP CONTRIBUTIONS

L.B. designed experiments, performed experiments, analyzed the data, performed some of the bioinformatics analyses, prepared the figures and wrote the manuscript. L.Br. performed experiments, analyzed the data and prepared the figures. M.D. performed experiments, analyzed the data and prepared the figures. C.D. performed most bioinformatics analyses and prepared the figures. L.D. performed bioinformatic analyses and prepared the figures. N.A. designed and performed experiments at the initiation of the project. L.G. performed single-cell and spatial transcriptomics experiments. M.F. and T.B. designed and performed the quantitative analysis of confocal microscopy images. B.E. performed the bioinformatics analysis of the spatial transcriptomics data. L.C. performed the spatial transcriptomics experiment. M.M., L.Be. and S.O. performed the influenza infection experiments. S.B. performed cell sorting for some experiments. C.S. and S.V.P. supervised the light-sheet microscopy staining, data acquisition, and data analyses. J.M.N. produced hashtag antibodies. M.G. supervised the influenza infection experiments. P.M. designed experiments, performed experiments, analyzed the data, prepared the figures, wrote the manuscript, acquired funding and supervised the study. All authors revised and approved the manuscript.

---

**Supplementary Figure 1.**
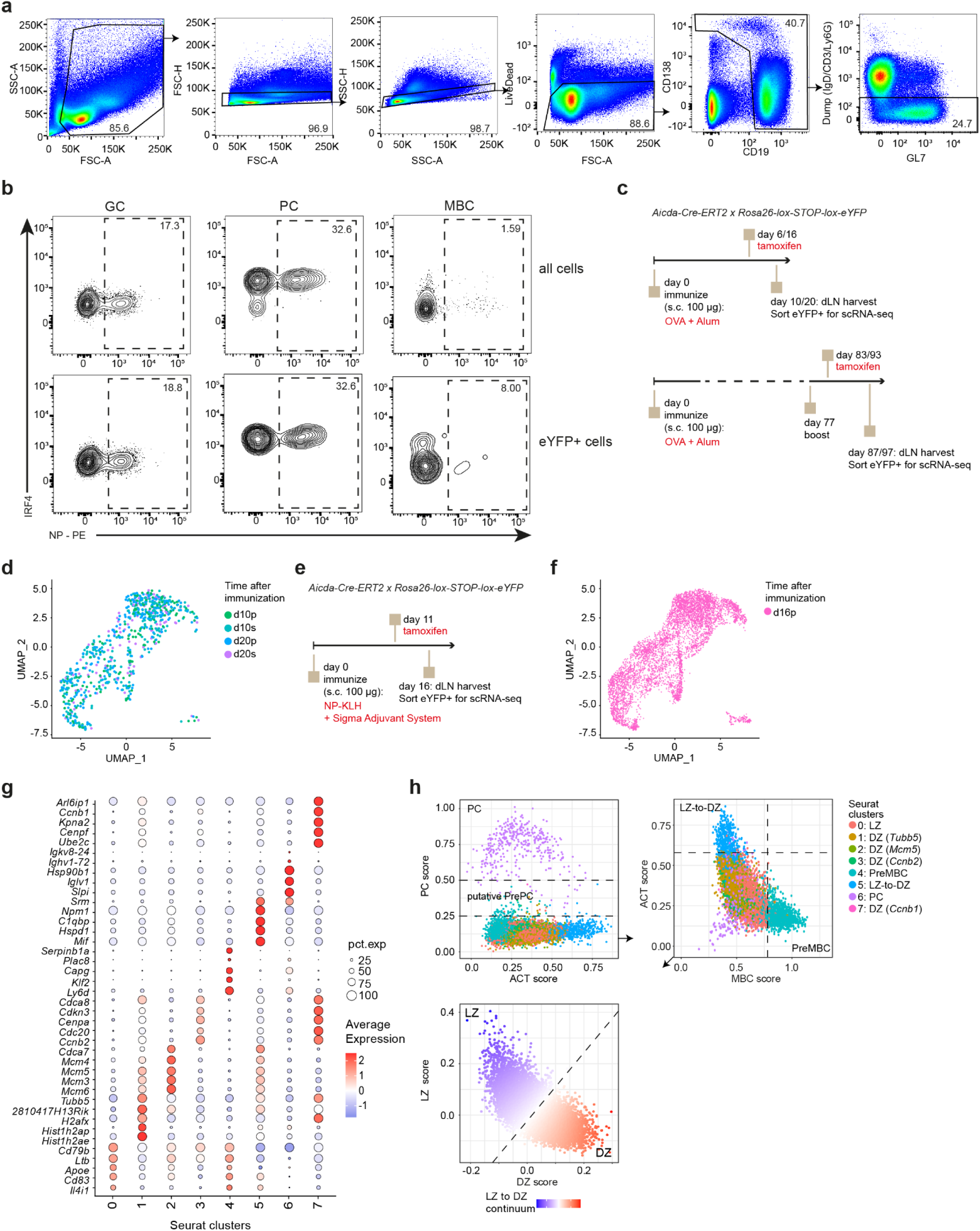
Single-cell RNA-seq analysis of fate-mapped GC B cells and recent GC emigrants identifies putative PrePC in draining LN after immunization (related to Figure 1). **a**, Flow cytometry gating strategy to identify live IgD^neg^ B cells from dLN of NP-KLH immunized mice. **b**, Flow cytometry gating of NP-specific cells among all (top panels) and eYFP^+^ (bottom panels) GC, PC and MBC (numbers indicate % of parent gate). **c**, Experimental design for the scRNA-seq analysis of GC and GC-derived cells in dLN after primary or secondary immunization with OVA antigen in Alum. **d**, UMAP representation of single-cell gene expression profiles of GC and GC-derived cells, colored by time after immunization (d10p: day 10 of primary response, d10s: day 10 of secondary response). **e**, Experimental design for the scRNA-seq analysis of GC and GC-derived cells in dLN after primary immunization with NP-KLH antigen in Sigma Adjuvant System. **f**, UMAP representation of single-cell gene expression profiles of GC and GC-derived cells, colored by time after immunization (d16p: day 16 of primary response. **g**, dot plot of the expression of top 5 markers of each unsupervised cluster, as indicated. The percentage of cells from a cluster expressing a marker is coded by the size of the circles, the average expression level by the color. **h**, Supervised gating strategy for the integrated scRNA-seq dataset based on gene expression scores for PC, activated (ACT), memory (MBC), DZ and LZ signature genes. Dashed lines represent the thresholds used to gate the cells in different annotations.

**Supplementary Figure 2.**
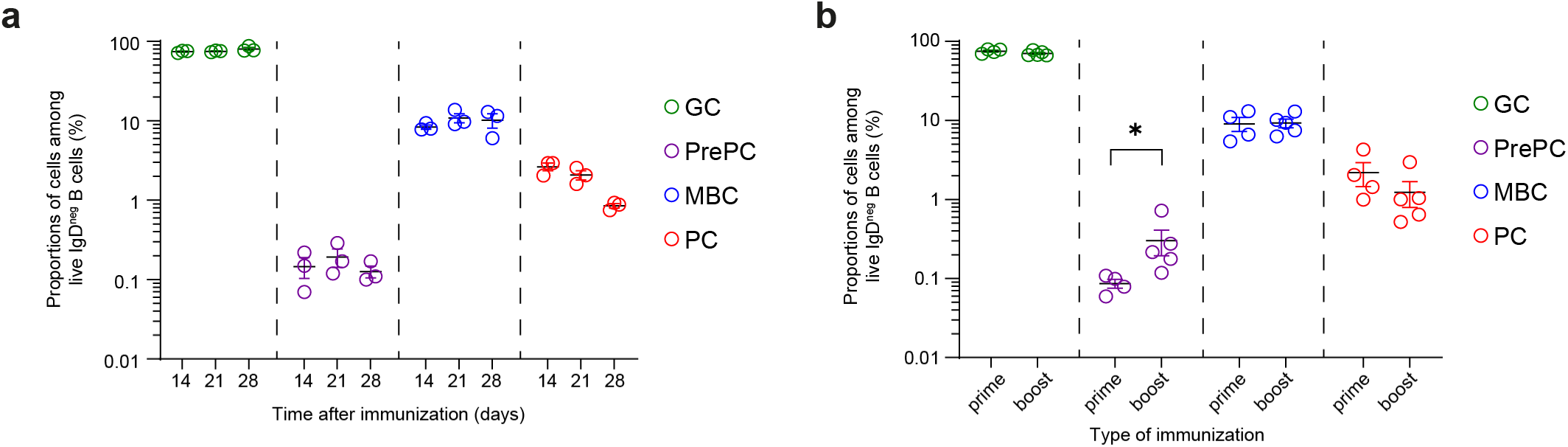
PrePC are generated in primary and secondary GC responses. **a**, Proportions of cells among dLN IgD^neg^ B cells from the indicated gates defined in Figure 2d in mice analyzed at different time points after primary immunization. **b**, Proportions of cells among dLN IgD^neg^ B cells from the indicated gates defined in Figure 2d in mice analyzed at day 15 after primary or secondary immunization. * p < 0.05 in Mann-Whitney non-parametric test.

**Supplementary Figure 3:**
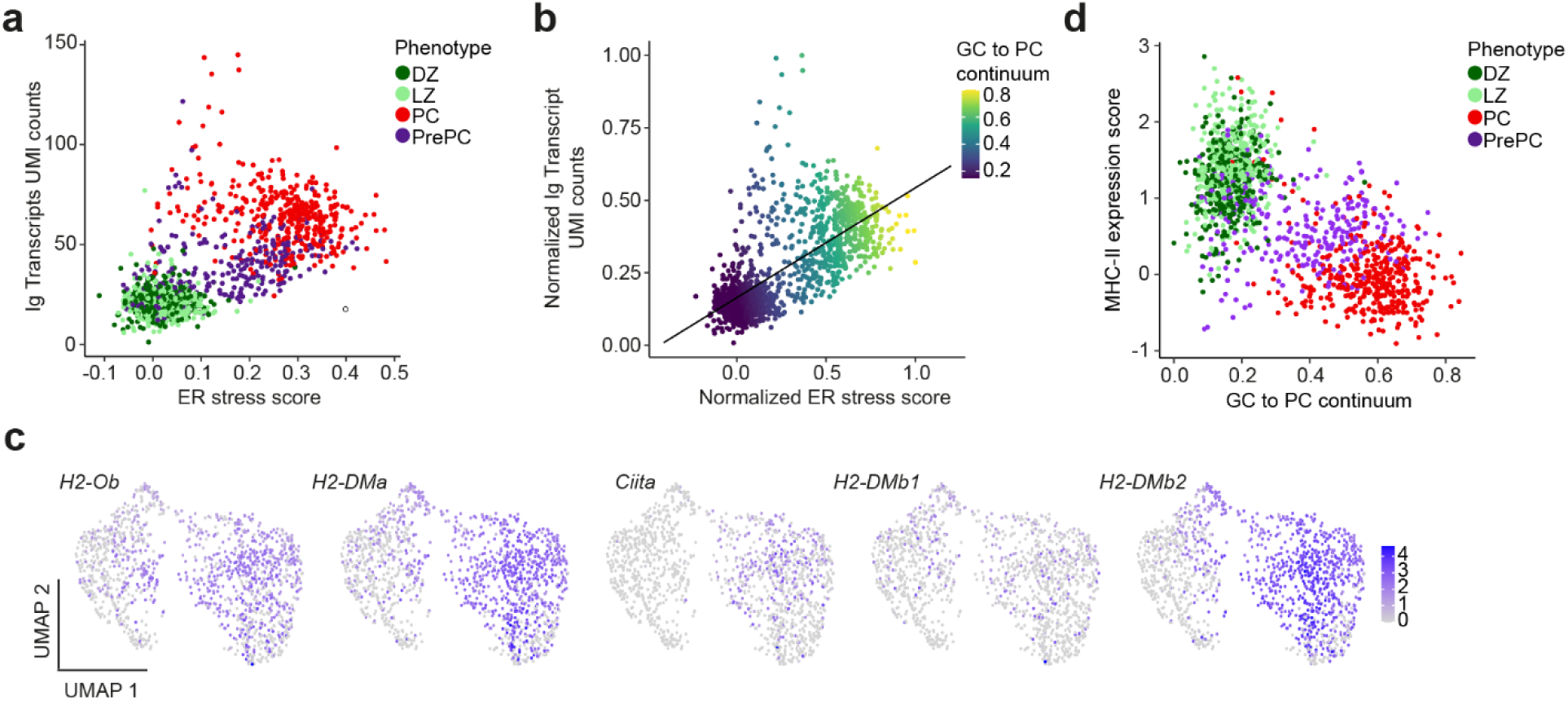
FACS-enriched PrePC cells from wild type immunized animals have an intermediate gene expression profile in the GC-to-PC differentiation continuum (related to Figure 3). **a**, Scatter plot representation of the ER stress score (x-axis) and the Ig transcripts UMI counts (y-axis) in cells colored by the FACS sorting phenotype. **b**, Scatter plot presented in a, overlayed with the regression line and colored by the GC to PC continuum score computed after projecting cells on the regression line and ranking. **c**, Feature plots showing the expression of the indicated MHC-II-related genes, laid out as in Figure 3c. **d**, Scatter plot representation of the GC to PC continuum score (x-axis) and the MHC-II gene expression score (y-axis) in cells colored by the FACS sorting phenotype.

